# The origin and maintenance of supergenes contributing to ecological adaptation in Atlantic herring

**DOI:** 10.1101/2023.10.23.562618

**Authors:** Minal Jamsandekar, Mafalda S. Ferreira, Mats E. Pettersson, Edward Farell, Brian W. Davis, Leif Andersson

## Abstract

Chromosomal inversions are associated with local adaptation in many species. However, questions regarding how they are formed, maintained and impact various other evolutionary processes remain elusive. Here, using a large genomic dataset of long-read and short-read sequencing, we ask these questions in one of the most abundant vertebrates on Earth, the Atlantic herring. This species has four megabase-sized inversions associated with ecological adaptation that correlate with water temperature. The *S* and *N* inversion alleles at these four loci dominate in the southern and northern parts, respectively, of the species distribution in the North Atlantic Ocean. By determining breakpoint coordinates of the four inversions and the structural variations surrounding them, we hypothesize that these inversions are formed by ectopic recombination between duplicated sequences immediately outside of the inversions. We show that these are old inversions (>1 MY), albeit formed after the split between Atlantic herring and its sister species, the Pacific herring. They are yet to reach mutation-flux equilibrium, but the large *Ne* of herring combined with the common occurrence of opposite homozygotes across the species distribution has allowed effective purifying selection to prevent accumulation of genetic load and repeats within the inversions.

## Introduction

Chromosomal inversions suppress recombination in the heterozygous state, facilitating the maintenance of different combination of alleles in tight linkage disequilibrium governing complex phenotypes, including the ones involved in local adaptation ^1–4^, reproductive strategies ^5^, life history traits ^6^, mimicry ^7^, and social behavior ^8^. Sets of alleles within the inversion are inherited together as a single unit in Mendelian segregation, and hence are also called supergenes^9^. Despite their evolutionary importance, the processes that lead to the origin, spread and maintenance of an inversion through time are often unclear because the evolution of inversion alleles is a dynamic process that changes over time and depends on the age, rate of gene flux, and effective population size (*N_e_*) of both inverted and non-inverted haplotypes ^10–13^. An inversion originates in a population as a single unit either by a recombination between near-identical inverted duplication sequences, a process known as nonallelic homologous recombination (NHAR), or by a repair mechanism of a single stranded break, known as Nonhomologous DNA End Joining (NEHJ) process ^14–16^. Such processes can only be understood by characterizing the breakpoint region, which is notoriously difficult to study as it is often present in the highly polymorphic part of the genome surrounded by complex structural variations (SVs) and repeats ^17,18^. Long-read sequencing makes it possible to uncover the complexity of breakpoints and shed light on the mechanisms forming inversions.

Immediately after its formation, a single inversion copy is vulnerable to the effects of random genetic drift, whereby it either can be lost or increase in frequency ^11^. If an inversion overlaps with co-adapted or beneficial allelic combinations, selection is likely to promote its maintenance and spread ^11^. However, suppressed recombination in heterozygotes can result in impaired purifying selection and consequent accumulation of deleterious mutations in the inversion region, which theoretical and empirical data have demonstrated to ultimately result in the degradation of the inversion through the process of Müller’s rachet ^6,7,19–23^. Interestingly, recent literature on vertebrate species supports the hypothesis that inversions can also evolve without pronounced accumulation of mutation load ^24–28^. The accumulation of mutation load depends on several factors, such as age and frequency of an inversion haplotype, as well as the *N_e_* of a species, with more efficient purifying selection in large populations ^10,12,13^. Furthermore, recombination may occur at low frequency in the heterozygotes, either through double cross-over or gene conversion, facilitating purifying selection and purging of deleterious mutations^29,30^. Our study uncovers such a process and thus contributes to the understanding of evolution of inversions.

In this study, we leverage the advancement in long-read sequencing technology with PacBio HiFi reads (average read length of 13.5 kb and accuracy above 99.8%) and use a large resequencing dataset to study four megabase-sized inversions on chromosomes 6, 12, 17, and 23 in the Atlantic herring (*Clupea harengus*) that are important for local adaptation ^31^. The variant haplotypes at these loci are denoted Southern (*S*) and Northern (*N*), owing to their respective predominance in the southern and northern parts of the species distribution range in the northern Atlantic Ocean, possessing warmer and colder waters, respectively ^31^. Atlantic herring is one of the most abundant vertebrates on Earth with an *N_e_* over a million and a census population size (*N_c_*) over a trillion, and has adapted to various ecological and environmental conditions such as variation in salinity, water temperature, light conditions, spawning seasons and food resources ^31,32^. The effect of random genetic drift should thus be minute with natural selection playing a dominant role in governing the evolution of genetic variation underlying ecological adaptation ^31^. Thus, Atlantic herring is an excellent model to explore the evolutionary history of supergenes associated with inversions in natural populations.

Here, we used whole genome PacBio HiFi data, combined with short-read Illumina data, from 12 Atlantic herring individuals, along with a previously generated high coverage re-sequencing dataset comprising 49 Atlantic herring and 30 Pacific herring (*Clupea pallasii*), the sister species ^31,33^ to shed light on (A) mechanisms of formation of inversion by finding breakpoint coordinates and structural variants (SVs) around the breakpoints, (B) the origin of inversions by describing its ancestral state and age using European sprat (*Sprattus sprattus*) as an outgroup species, (C) evolutionary history of inversions by phylogeny, (D) effects of suppressed recombination by analyzing patterns of variation, differentiation, linkage disequilibrium, mutation load, and gene flux (genetic exchange between inversion haplotypes).

## RESULTS

### Samples and genome assemblies

The analysis of short read as well as long read data showed that the six Celtic Sea samples (CS2, CS4, CS5, CS7, CS8, and CS10) were homozygous for the Southern (*S*) allele for all four inversions (Supplementary Figs. 1, 2). Among the six Baltic Sea samples (BS1-BS6), four were homozygous for all Northern (*N*) alleles while two samples, BS2 and BS5, were heterozygous for the Chr23 and 17 inversions, respectively (Supplementary Figs. 1, 2). All samples were sequenced using PacBio HiFi with coverage ranging from 23x to 30x. We produced 24 haploid *de novo* genome assemblies from the 12 herring samples using hifiasm ^36^. The two haploid assemblies per individual were denoted hap1 and hap2. All assemblies were of high quality, where genome size ranged from 743 to 792 Mb, the contiguity measured by N50 ranged from 452 to 737 kb and BUSCO score was above 90%, indicating that PacBio assemblies contained more than 90% of conserved vertebrate genes (Table 1, Supplementary Table 1). Notably, hap1 assemblies had larger genome size and more contigs as compared to the hap2 assemblies. The unequal genome size for two haplotype assemblies suggests that a minor fraction of heterozygous sequences might not be accurately phased; while the positive correlation between genome size and number of contigs suggests that a small fraction of the genome is fragmented into multiple contigs.

**Table 1:** Genome statistics for all CS10 and BS3 de novo genome assemblies. Hap1 and hap2 are the two haplotype genomes and their statistics are shown on the top and bottom row for each sample, respectively.

| Samples | Genome size (Mb) | Genome completeness (%) | No. contigs | N50 (kb) | N's per 100 kb |
| --- | --- | --- | --- | --- | --- |
| **CS10** | 773.1 | 90.7 | 3997 | 452.8 | 0 |
|  | 756.6 | 90.3 | 3228 | 472.4 | 0 |
| **BS3** | 779.2 | 92.4 | 3331 | 680.2 | 0 |
|  | 758.5 | 92.1 | 2725 | 661.8 | 0 |
| **Reference** | 725.7 | 87.4 | 26 | 30,022.48 | 110.78 |
|  | 786.3 | 93.3 | 1697 | 29,845.74 | 118.43 |

We compared the quality of PacBio assemblies with that of the reference assembly of Atlantic herring (Ch_v2.0.2) and found that PacBio assemblies were of similar quality for genome size and BUSCO scores (Table 1). Although the total size of the reference assembly is 786 Mb, only 726 Mb is scaffolded in chromosomes (*n* = 26) and the remaining 61 Mb is present as unplaced scaffolds (*n* = 1,697), i.e., fragments that could not be assigned to a chromosome. This material likely includes unresolved haplotypes, which is supported by the fact that all novel PacBio assemblies were above 726 Mb, indicating that these assemblies have higher portions of the heterozygous alleles resolved into separate contigs than its reference counterpart, which would be expected due to the improvement in accuracy provided by the HiFi technology.

### Characterization of inversion breakpoints on chromosomes 6, 12, 17, and 23

Alignment of HiFi reads from the alternate inversion allele spanning the inversion breakpoint to the reference assembly is expected to show a particular pattern where the reads get split into two parts, one part aligns outside the inversion and the other part aligns inside the inversion at the opposite end in reverse orientation. We manually inspected our data for such a pattern using IGV and Ribbon and revealed the inversion breakpoint in all samples for all four inversions (six samples for each inversion). We found similar breakpoint co-ordinates for all samples in each inversion (Fig. 1, Supplementary Table 2), suggesting that these inversions have originated just once, stemming from a one-time break in the chromosome, and have not reoccurred multiple times using the same breakpoint regions; a pattern observed in some other species ^78,79^. Chr6 slightly deviated from this common observation as we find that the distal breakpoint for one of the samples was 500 kb further along the chromosome (Supplementary Fig. 3, Supplementary Table 2). This could be due to either a different distal breakpoint or a secondary inversion, but more samples are needed to confirm these possibilities. We re-evaluated the breakpoints obtained in our previous study ^31^ for Chr6 and Chr17 inversions and found 1-3 kb shifts at two positions (Supplementary Table 2). Fig. 1 reports the breakpoint co-ordinates that occurred more frequently in the examined samples, which were Chr6:22,282,765-24,868,582, Chr12:17,826,318-25,603,093, Chr17:25,802,2019-27,568,510, Chr23:16,225,343-17,604,279. None of these breakpoints disrupted any coding sequence (Fig 1; the list of genes around the breakpoints is provided in Supplementary Table 3). The inversions were further confirmed by the alignments of *N* and *S* allele scaffolds constructed using PacBio contigs and optical mapping data (Fig. 2). The breakpoint coordinates on *N* and *S* allele scaffolds were determined by noting the coordinates where the scaffolds change orientation in the sequence alignment dot plot (Supplementary Table 4).

**Fig. 1:**
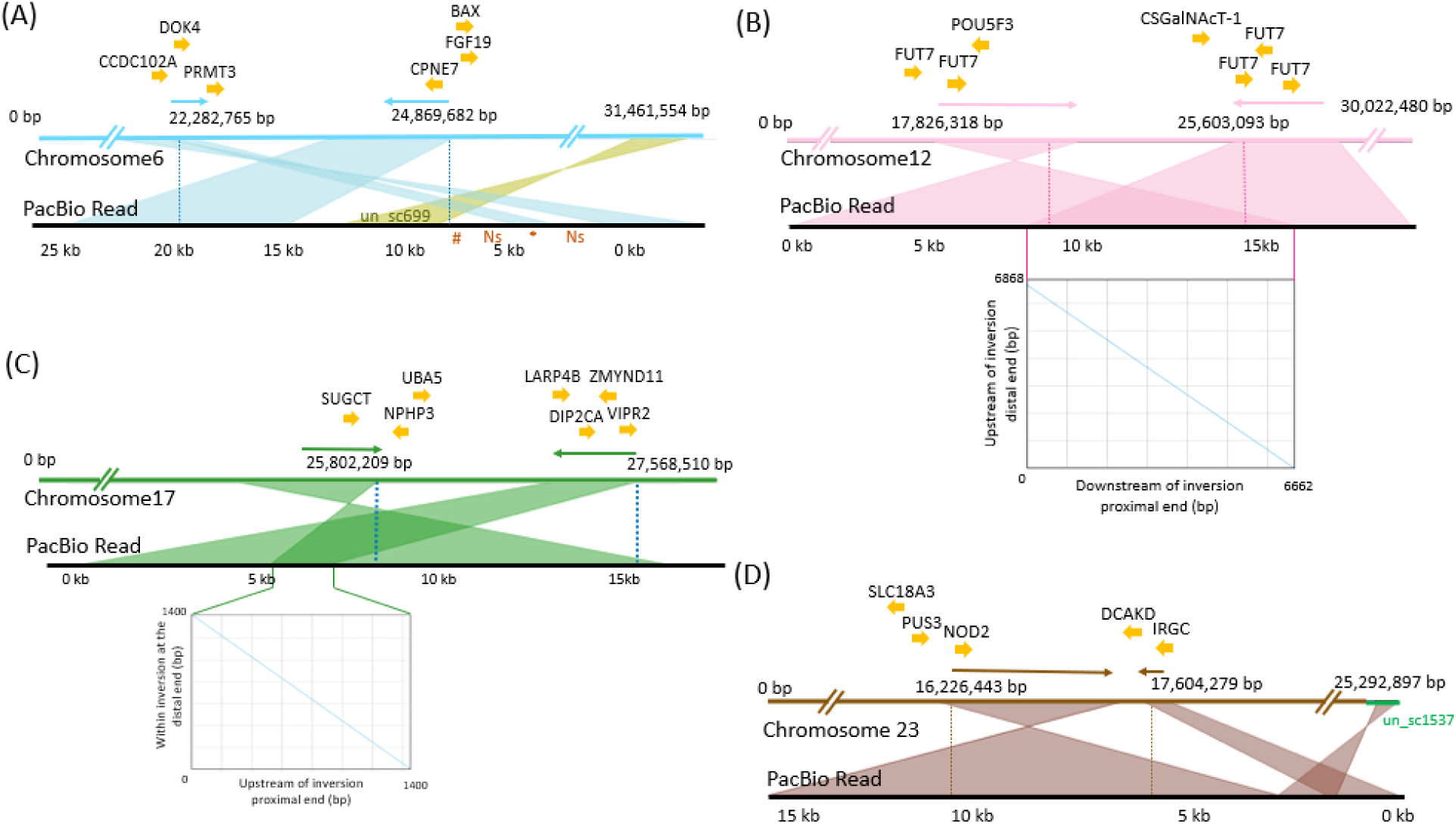
Single PacBio reads spanning proximal and distal breakpoints on the reference assembly. The orientation of the HiFi reads are indicated with arrows above the reference sequence. The inversion breakpoints are shown with dotted lines. Genes at the breakpoints are indicated above the reference sequence. **(A) Chromosome 6** (modified from ^31^). **(B) Chromosome 12.** *FUT7*, Fucosyl transferase 7; *POU5F3*, POU domain, class 5, transcription factor 3; *CSGalN-Act1a*, chondroitin sulfate N-acetyl galactosaminyl transferase 1a. **(C) Chromosome 17** (modified from ^31^). **(D) Chromosome 23**. A part of the HiFi read is mapped to the unplaced_scaffold (un_sc) 1537 on the reference, which is shown in green here. *SLC18A*, probable vesicular acetylcholine transporter-A; *PUS3*, pseudouridylate synthase 3; *NOD2*, nucleotide-binding oligomerization domain containing 2; *IRGC*, interferon-inducible GTPase 5- like; *DCAKD*, dephospho-CoA kinase domain containing protein. The dot plots below the read alignments for chromosomes 12 and 17 compare two copies of the duplication present at the breakpoint, which are also indicated by an overlap of the read at proximal and distal ends of the inversions. The length of the duplication on chromosomes 12 and 17 are 8 kb and 3 kb, respectively.

**Fig. 2:**
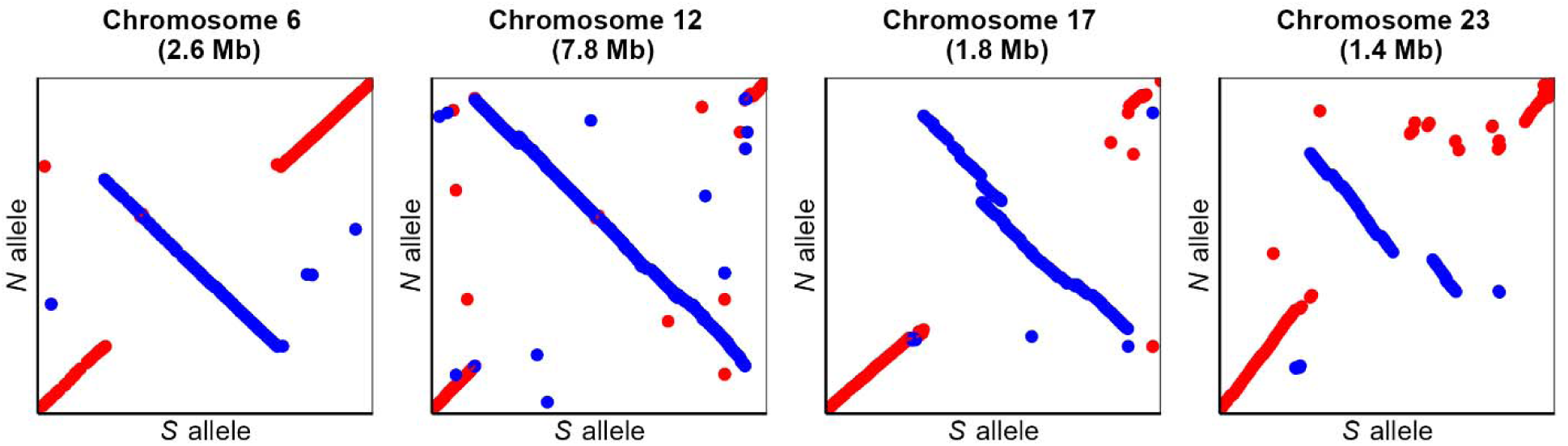
Sequence alignment of *N* and *S* alleles for all four inversions. *N* and *S* alleles are represented by the CS10_hap1 and BS3_hap2 inversion scaffolds, respectively.

### Identification of ancestral haplotypes using European sprat as an outgroup

An inversion allele will initially have lower nucleotide diversity (*π*) than its ancestral counterpart, and thus *π* estimates can be used to indicate which allele is derived and which is ancestral if the inversion is a relatively recent event ^13^. However, if the inversion is old, similar to the coalescence time for neutral alleles or older, *π* will be similar for the two alleles. Further, if gene flux occurs between alleles (see below), diversity may be homogenized between inversion haplotypes and thus differences in *π* estimations may be abolished. In fact, *π* estimates for the *N* and *S* haplotypes are similar for all four inversions ^31^. We therefore decided to use the European sprat (*Sprattus sprattus*) as an appropriate outgroup species ^50,51^ to determine which inversion haplotype represents the derived state. Based on the linear orientation of the alignment of sprat contigs to the *N* and *S* alleles, we concluded that *S* is the ancestral haplotype for the inversions on Chr6 and 12; while *N* is the ancestral haplotype for the inversions on Chr17 and 23 (Fig. 3). However, results for Chr23 should be treated with caution as the alignment was fragmented.

**Fig. 3:**
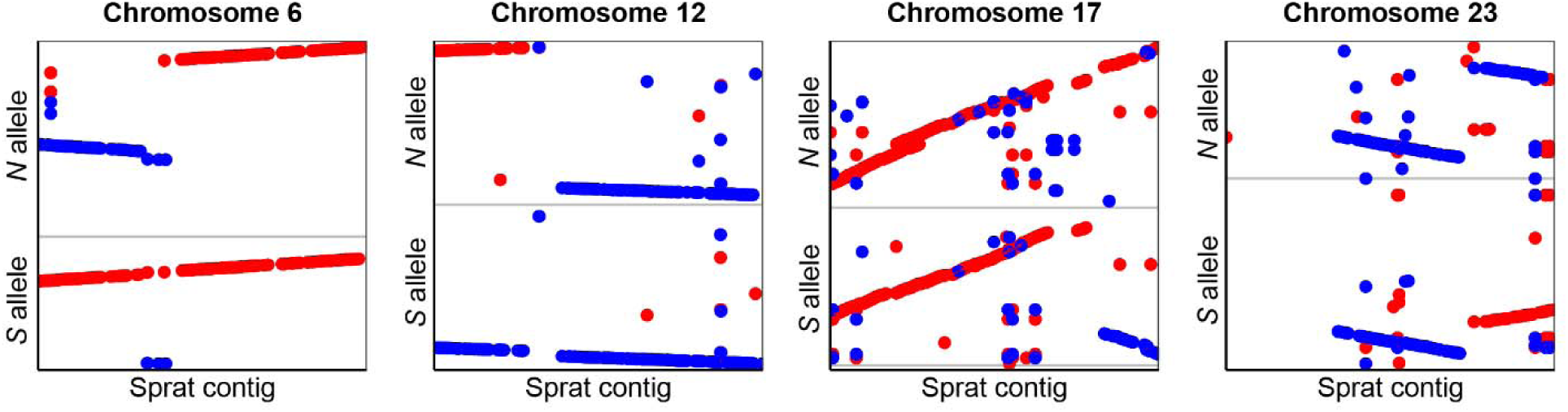
Sequence alignment of Atlantic herring inversion alleles and contigs from the European sprat assembly spanning the inversion breakpoints for all four inversions.

### Structural variations at the breakpoint regions

Leveraging the long PacBio contigs spanning the inversion breakpoints, we studied structural variants (SVs) and repeats surrounding the inversion breakpoints in each haplotype, which could have played a role in the formation of the inversions. The sequence alignments of *N* and *S* alleles near the breakpoints indicated that the breakpoints for all four inversions were flanked by inverted duplications ranging from 8-60 kb in size and contained one or no gene (Fig. 4 and Supplementary Fig. 4). Such duplicated arrangements can facilitate ectopic recombination resulting in the formation of inversions. In addition, other types of SVs like indels, palindromes, duplications were also enriched near the breakpoints (Supplementary Table 5).

**Fig. 4:**
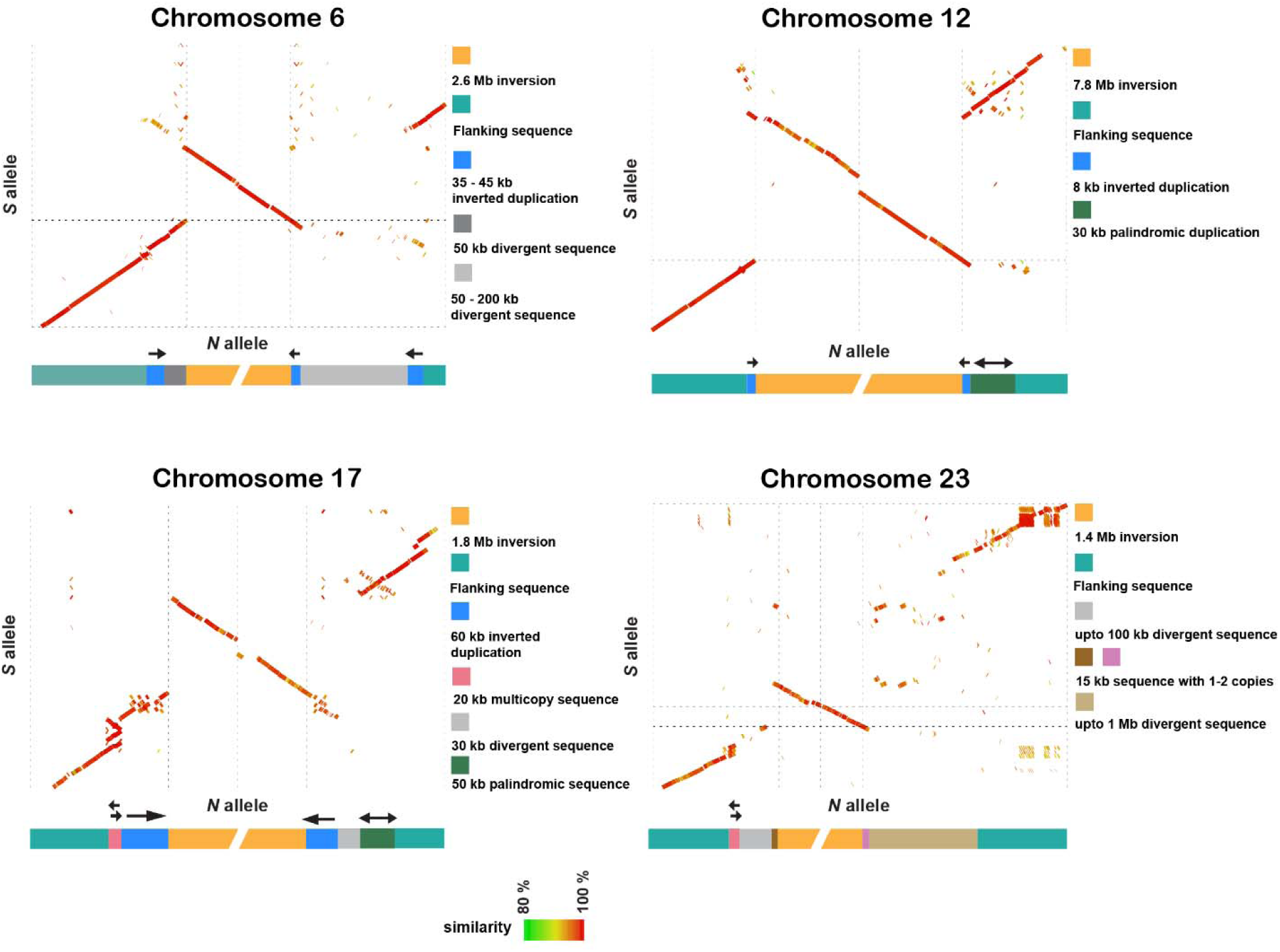
Sequence alignments and models at the breakpoint regions for the *N* and *S* inversion scaffolds. The green to red color gradient in dot plots represents percent sequence similarity from 80 to 100. The models representing SVs surrounding the inversions are below the dot plots. Chr6: 400 kb sequence includes 300 kb outside inversion and 100 kb inside inversion for both proximal and distal regions. Chr12: 200 kb sequence includes 100 kb outside inversion and 100 kb inside inversion for both proximal and distal regions. Chr17: 400 kb sequence includes 300 kb outside inversion and 100 kb inside inversion for both proximal and distal regions. Chr23: 400 kb proximal. 600 kb distal sequence includes 500 kb outside inversion and 100 kb inside inversion. The plots are based on the CS10_hap1 and BS3_hap2 genome assemblies.

We also studied SVs near and inside inversion haplotypes using genome graphs (Supplementary Fig. 5), which corroborated the complexity of breakpoint regions already apparent in the dot plot analysis (Figure 4). For instance, distal breakpoints of all inversions were divergent among individuals, revealing the existence of non-shared structural variants. In particular, the Chr17 and Chr23 breakpoints were the most complex. The distal breakpoint of Chr17 coincided with a telomeric sequence that varies in length (0-300 kb) outside the breakpoint and that is misaligned in the genome graph. The Chr23 inversion breakpoints were the most divergent across individuals, revealing the existence of a long breakpoint region with many structural variants. This complexity suggests that, after the formation of an inversion, there could be an accumulation of structural variants around the breakpoints, in this case, not associated with any particular inversion allele and that may be evolving neutrally. The genome graphs (Supplementary Fig. 5) also revealed the existence of structural variants inside inversions, in particular in Chr12, 17 and 23, while Chr6 alignments revealed higher similarity among haplotypes.

### Origin of inversion haplotypes

We used short-read sequencing datasets of Atlantic and Pacific herring individuals to study the origin and subsequent evolution of the inversions. We first studied the genome-wide evolutionary history of the two *Clupea* sister species, Atlantic and Pacific herring, by either (A) using a concatenate alignment of 15,471 genes (∼114 Mb with no missing data) including the European sprat to generate a rooted tree, or (B) by using a longer ∼346 Mb genome-wide alignment with no missing data containing only *Clupea* individuals, to more confidently infer intraspecific relationships. Atlantic and Pacific herring formed well-supported monophyletic sister clades (Fig. 5A-B and Supplementary Fig. 6). Maximum likelihood trees of all four inversion haplotypes using data from multiple herring populations revealed a similar evolutionary history as the genome-wide species tree (Fig. 5 and Supplementary Fig. 7). In rooted and unrooted trees, we found a split of all Atlantic herring individuals from the Pacific herring, followed by a split between reciprocal homozygotes of each inversion allele, with heterozygotes placed between these two clusters (Fig. 5C and Supplementary Fig. 7), suggesting that the inversions originated after the split between Atlantic and Pacific herring. The *S* cluster is constituted by all individuals originating from Britain and Ireland and part of the North Sea individuals in all four trees, whereas the *N* cluster is mostly constituted by Baltic Sea, Norwegian and Canadian herring. The coincidence of phylogenetic relationship between *N* and *S* alleles and geographic distribution of the individuals is in line with previous results that suggest that the *S* alleles at each inversion tend to occur at high frequency in warmer waters particularly around Britain and Ireland, whereas *N* alleles occur at high frequency in colder waters in the north ^31,80^.

**Fig. 5:**
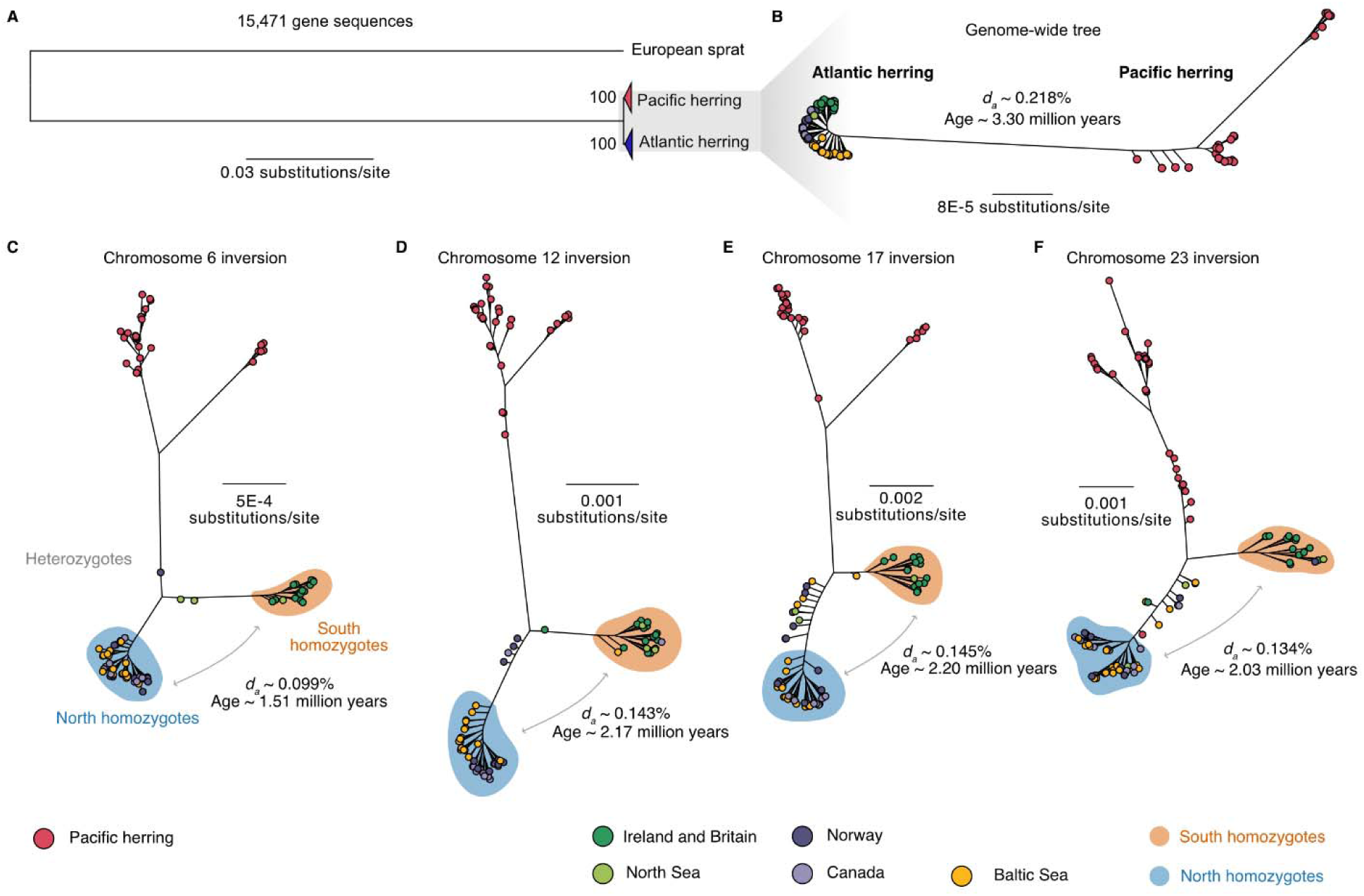
The evolutionary history of chromosomal rearrangements in Atlantic herring. (A) Maximum likelihood tree of 15,471 concatenated gene sequences from 30 Pacific and 61 Atlantic herring, using the European sprat as an outgroup; **(B)** Maximum likelihood tree of 345,966,161 concatenated genome-wide positions with no missing data using the same 91 individuals; **(C-F)** Maximum likelihood trees of the inversion regions (alignments allowing 50% missing data). Estimated net nucleotide divergence (*d_a_*) and divergence times in million years between Atlantic and Pacific herring **(B)** and between *N* and *S* homozygotes **(C-F)** are indicated. In trees **(B-F)**, Atlantic herring individuals are color coded depending on the population of origin.

We used net nucleotide diversity (*d_a_*) between the *N* and *S* homozygotes to estimate divergence among inversion haplotypes, relative to the divergence of Atlantic and Pacific herring (Fig. 5). Divergence times ranged from 1.51 million of years (MY) (Chr 6) to 2.20 MY (Chr 17), which are more recent divergences than the one estimated for Atlantic and Pacific herring in our dataset (3.30 MY). Given that *d_xy_* values between *N* and *S* alleles were lower but close to *d_xy_* between Atlantic and Pacific herring across inversion regions (Supplementary Fig. 8A), and given the possibility of recombination among inversion haplotypes (Supplementary Figs. 5, 2 and results below) ^29^, it is possible that our estimated divergence times are underestimations, suggesting that the inversions are old polymorphisms that could have originated shortly after the split between the two *Clupea* species.

### The evolutionary history of inversion haplotypes

We explored the evolutionary history of the four inversions using homozygous individuals from the Baltic and Celtic Sea (total *N*=35; see Supplementary Fig. 2) and calculated sequence differentiation (*F*_ST_), nucleotide diversity (*π*), and linkage disequilibrium (LD) measured as *R*^2^ across the inversions and their flanking region (Fig. 6, Supplementary Fig. 11). The inversion regions showed strong differentiation between *N* and *S* homozygotes (high *F*_ST_) which is in sharp contrast with the flanking regions (low *F*_ST_). An exception to this is a region proximal to the Chr17 inversion breakpoint. A careful inspection of our PacBio data showed that this is not part of the inversion and must be a sequence polymorphism in very strong LD with the inversion polymorphism. The LD across all four inversions was strong, particularly for Chr6 and Chr12 inversions. Nucleotide diversity for all four inversions showed significant differences between haplotypes and genome-wide averages in certain cases, but *π* values of all inversions are within the genome-wide distribution of *π* (Fig. 6) The nucleotide diversity of inversion alleles representing the derived state is not lower than for those representing the ancestral state (Fig. 6), as only the Chr 12 inversion showed a significantly reduced diversity in the derived haplotype (*P* < 0.001 for Chr12) as expected, while Chr17 and Chr23 inversions showed higher diversity in the derived haplotype (*P* << 0.001 for Chr17 and *P* < 0.01 for Chr23) (Fig. 6). These results are consistent with the old age of the inversion polymorphisms (Fig. 5), exceeding the coalescence time for neutral alleles in Atlantic herring.

**Fig. 6:**
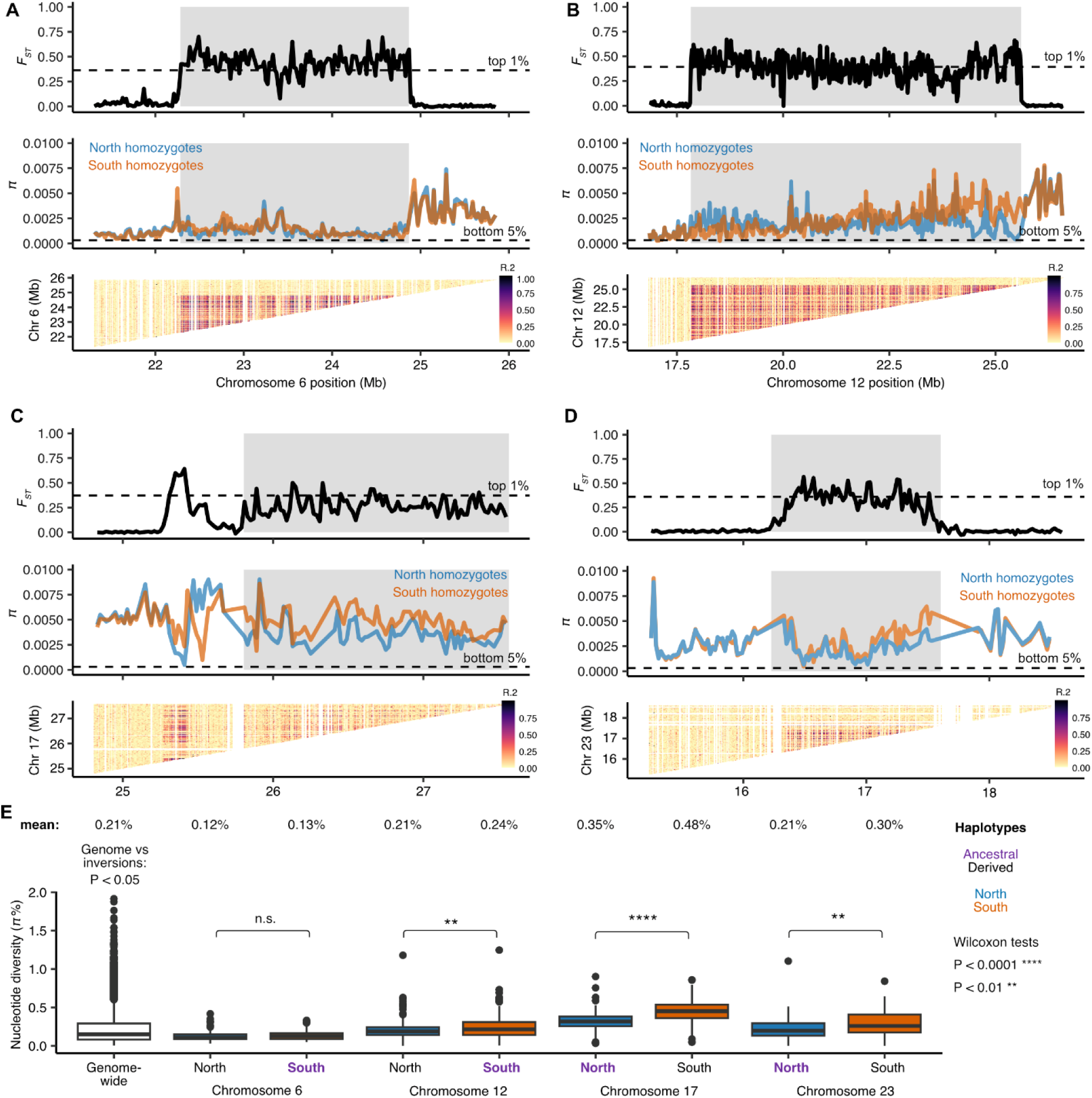
Differentiation, diversity and recombination patterns in inversion regions in Atlantic herring on (A) Chr6, (B) Chr12, (C) Chr17 and (D) Chr23. Distribution of *F*_ST_ and nucleotide diversity (*π*) for *N* and *S* homozygotes are displayed in sliding windows of 20 kb across inversion regions (shaded gray boxes). Linkage disequilibrium is represented as *R*^2^ among genotypes in the inversion region. **(E)** Comparison of the distribution of nucleotide diversity (*π*) calculated in sliding windows of 20 kb for each inversion haplotype and genome-wide. Averages are displayed above the boxplots and ancestral alleles are highlighted in purple.

Suppression of recombination between inversion haplotypes is expected to accumulate deleterious mutations and transposable elements (TEs) due to impaired purifying selection, as the inversion haplotypes have a reduced *Ne* compared with the rest of the genome ^22^. To test this, we compared the number of nonsynonymous substitutions per non-synonymous site (*d_N_*) to the number of synonymous substitutions per synonymous site (*d_S_*), or *d_N_/d_S_* and site frequency spectrum (SFS) of non-synonymous and synonymous mutations for genes within the inversions to the genome average, using the European sprat as an outgroup species. We found no significant difference in *d_N_/d_S_* for any inversion allele and the genome-wide distribution (*P* > 0.05, two-sided *t*-test, Fig. 7A). Further, the SFS of *N* and *S* homozygotes were similar to each other (Fig. 7B) and to that of the genome-wide estimate (Supplementary Fig. 9), where polymorphic synonymous positions are always the most abundant class, suggesting that low frequency non-synonymous mutations are being effectively purged from inversion haplotypes. Furthermore, the observed derived alleles at high frequencies are candidate mutations for being under positive selection. Further, we compared the *d_N_/d_S_* ratio for the *N* and *S* alleles at each locus in an attempt to find genes that may show accelerated protein evolution as part of the evolution of these adaptive haplotypes. However, the ratios were remarkably similar in pairwise comparisons with only a few genes showing a minor difference in *d_N_/d_S_* (Supplementary Fig. 12).

**Fig. 7:**
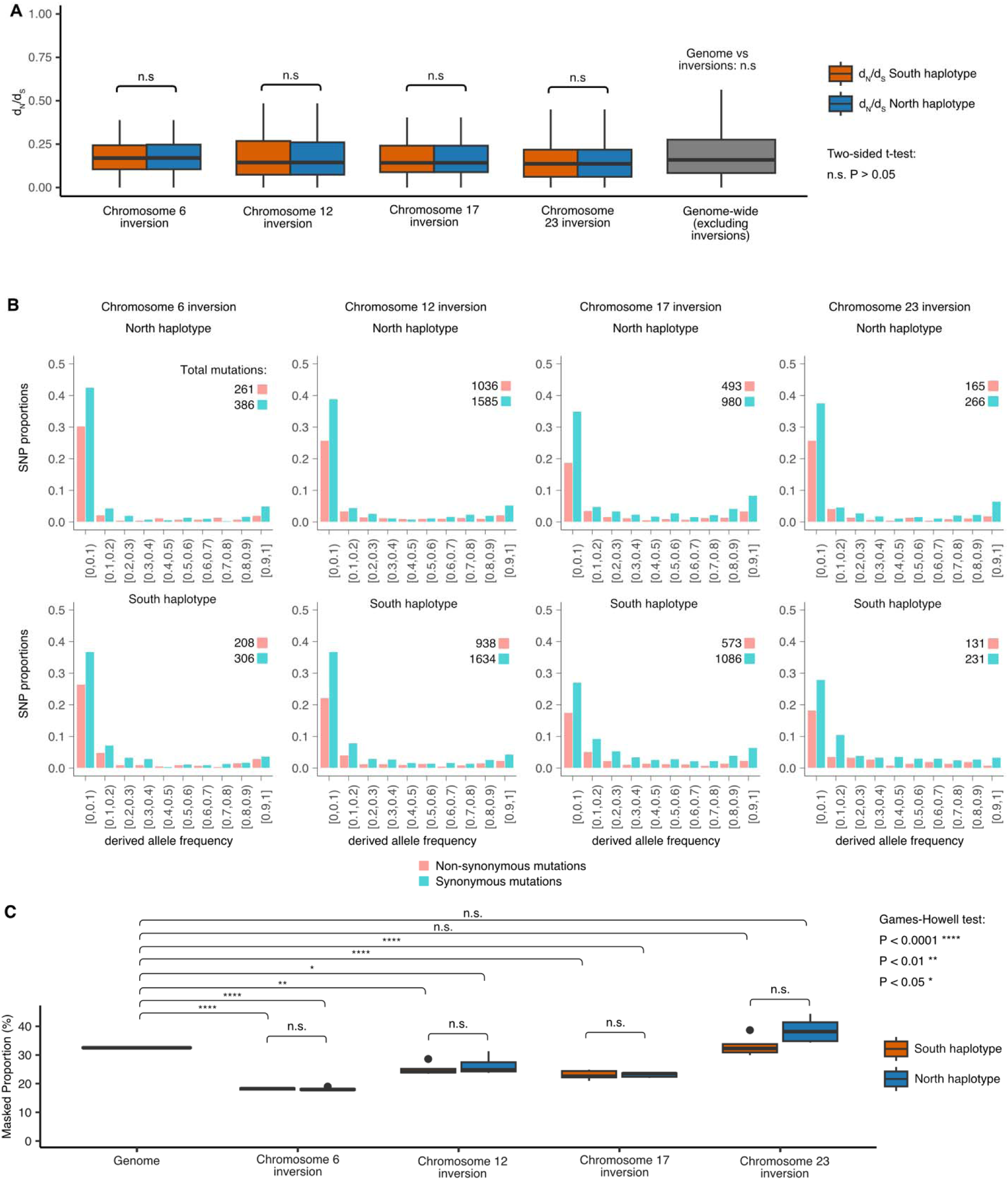
Lack of mutational load in *N* and *S* inversion haplotypes. (A) Distribution of *d_N_/d_S_*ratios for *S* and *N* alleles of genes overlapping with the inversions on chromosomes 6, 12, 17 and 23 compared with the genome-wide *d_N_/d_S_* distribution generated with all 61 individuals in dataset (excluding genes inside inversions). A two-sided *t*-test revealed no significant differences in the distribution of *d_N_/d_S_* values between *N* and *S* alleles, or between inversion haplotypes and the genome. **(B)** Site frequency spectra of derived non-synonymous and synonymous mutations for each inversion haplotype. The total number of mutations in each category is displayed above the graph. **(C)** Proportion of transposable elements for the entire genome and for each inversion haplotype. A Games-Howell non-parametric test was performed to test differences between proportion of TEs between each allele and the genome (n.s. = non-significant).

Finally, we compared TE abundance between *N* and *S* haplotypes, as a proxy of mutational load, which revealed non-significant difference between haplotypes and a lower TE content in Chr6, Chr12 and Chr17 inversions compared to the rest of the genome (Fig. 7C). Taken together, the data on *d_N_*/*d_S_* and on TE content did not indicate increased genetic load for alleles at any of the four inversion polymorphisms strongly associated with ecological adaptation.

### Evidence of allelic exchange between inversion haplotypes

To visualize genetic exchange between inversion haplotypes, we constructed a deltaAlleleFrequency’(dAF’) metric that measures the degree of allele sharing between haplotypes (see Methods). dAF’ = 1.0 means that there is a maximum dAF given the frequencies of sequence variants among haplotypes, while dAF’ = 0 means that sequence variants have the same frequencies among the two haplotypes. All sequence variants within an inversion will show dAF’ = 1 if there has been no recombination between haplotypes and the same mutation has not occurred on both haplotypes. This analysis documents extensive allele sharing at all four loci, and in particular for Chr12 and 17 (Fig. 8), because if there had been no recombination between haplotypes, all SNPs within the inversion would have been colored blue in Fig. 8. The result is consistent with our previous analysis of allele sharing for the Chr12 inversion ^41^. The region between Chr12: 23.0-23.5 Mb, with a particularly high incidence of sequence variants with low dAF’ values correspond to an interval where we have noted evidence for genetic recombination between the *N* and *S* alleles ^41^, where we see a drop in *F*_ST_ between haplotypes (Fig. 6) and an excess of heterozygous genotypes in Baltic Sea individuals (Supplementary Fig. 2). This analysis also confirms the extreme sequence divergence between *N* and *S* homozygotes in a flanking region outside the inversion for the Chr17 inversion. Extremely differentiated SNPs (dAF > 0.95) were not enriched for non-synonymous mutations (Supplementary Table 6, list of genes with non-synonymous mutations is in Supplementary Table 7).

**Fig. 8:**
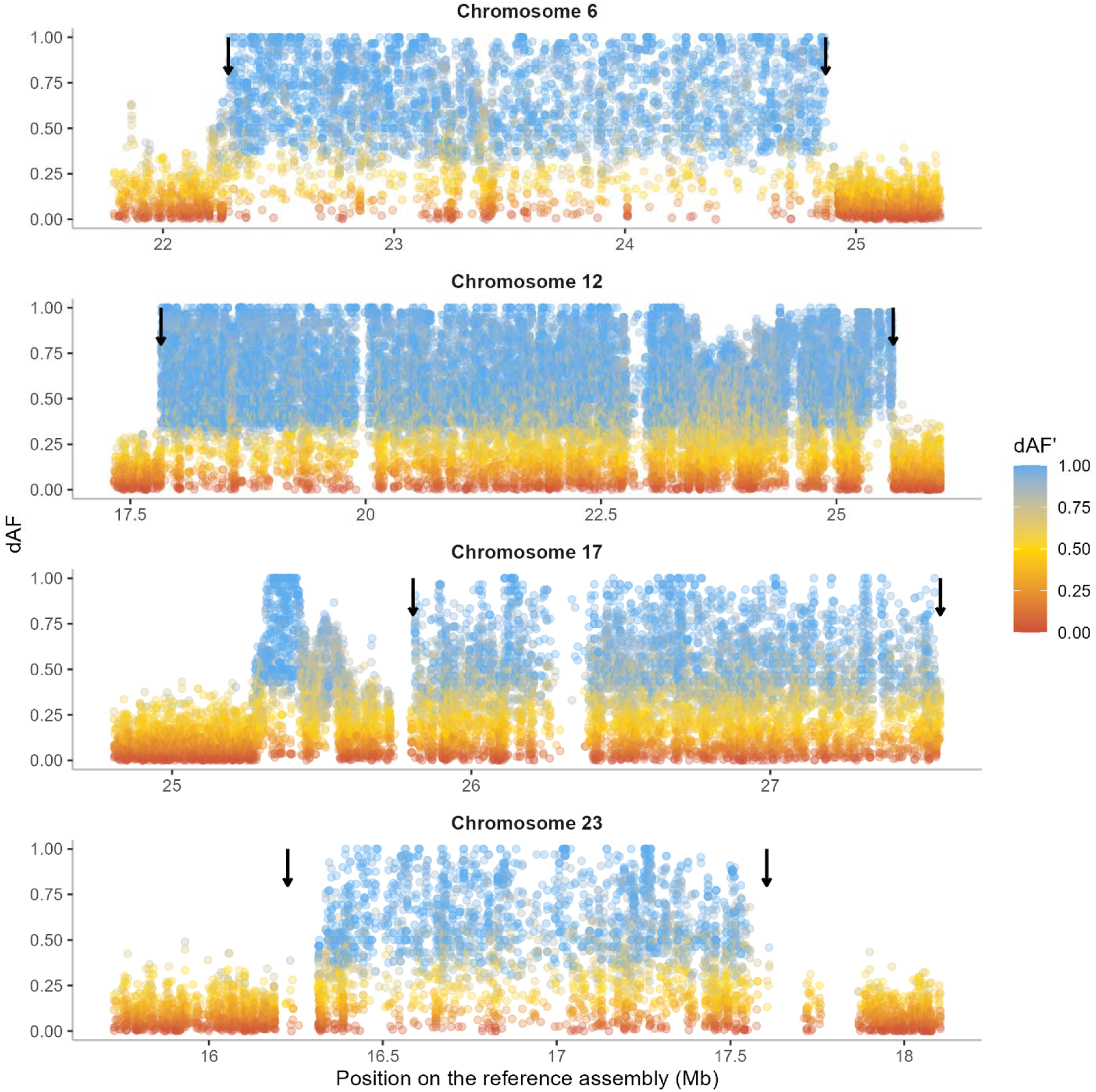
dAF between *N* and *S* populations homozygotes for four inversions, colored by dAF’. Arrows indicate the inversion breakpoints. The blank regions on chromosomes 12, 17, and 23 represent regions where SNPs are not called due to complexity of the genomic regions with many repeats and indels.

## Discussion

In this study, we were able to leverage the power of long-read sequencing to confirm the presence of four large inversions in Atlantic herring that all show strong differentiation between populations from the northern versus southern part of the species distribution ^31^. By comparing our assemblies to long-read data for an outgroup species, European sprat, we were able to determine that the *S* arrangement is ancestral at Chr6 and Chr12, whereas *N* is ancestral at Chr17 and Chr23. This is different than our previous prediction for Chr17 that was based on the comparison of duplicated sequence ^31^. We also used extensive re-sequencing data from Atlantic herring and its sister species, the Pacific herring, to study the evolutionary history of these inversion polymorphisms. Our phylogenetic analysis shows that these four inversion polymorphisms have been maintained for more than a million years but they most likely all occurred subsequent to the split between Atlantic herring and its sister species, the Pacific herring. Furthermore, we also studied patterns of diversity, differentiation, recombination, mutation load accumulation and genetic exchange in the inversion regions to get insights into the evolutionary history of the inversions after their formation. Our results reveal no indication of accumulation of mutation load, despite high differentiation and strong suppressed recombination in the region. The signatures of genetic exchange between inversions and their high frequency in natural populations suggests inversions have been maintained for a long evolutionary period by divergent selection.

### Inverted duplications present near inversion breakpoints

Here we characterized the breakpoint regions in detail, shedding new light onto the role of structural variation in the origin of the inversions and their subsequent evolution. We found very similar chromosomal breakpoints in all haplotypes (Supplementary Table 2) suggesting that each of the four inversions has a single origin. Phylogenetic trees of the inversion haplotypes also support this result, since individuals cluster by their genotype at the inversion (*NN* or *SS*), rather than geographic location (Fig. 5). The data support a single origin of the inversions after the split between Atlantic and Pacific herring. The inversion breakpoints of Chr6, 12, and 17 were flanked by inverted duplications (Fig. 4), setting the stage for nonallelic homologous recombination (NAHR) (also referred to as ectopic recombination) leading to the formation of inversion. This mechanism of inversion formation is commonly accepted ^16^ and reported in a few species of *Drosophila* and eutherian mammals ^25,81–83^.

As we find no inverted duplications for inversion on Chr23, it is possible that it has been formed by the alternative mechanism of double strand staggered breaks. This mechanism is argued to be the most common mechanism in invertebrate species, where a single stranded break is repaired by Nonhomologous DNA End Joining (NHEJ) and results in inversion accompanied by duplication ^14,15^. Nevertheless, NAHR seems to be the most common mechanism in vertebrates ^25,84^ and indeed our data supports a prevalent role of NAHR in the formation of at least three of the four inversions in Atlantic herring. The presence of flanking inverted duplication sequences increases the probability of recurrent inversions, an event termed as “inversion toggling”, by breakpoint reusage ^85^. All the breakpoints were surrounded by repeated, palindromic, and divergent sequences (Fig. 4 and Supplementary Fig. 5), which could have been formed by a gene conversion process using the inverted duplicates flanking the inversion breakpoints. Such process can further facilitate the formation of insertions and deletions, during which, double stranded breaks, strand extension, and rejoining create even more duplicated sequences ^81^. Notably, most of these SVs outside of inversions occur in non-genic regions. Presence of such divergent sequences around breakpoint might be responsible for restricting the gene flow at the breakpoints and thus maintaining the diversity among haplotypes, as peaks of divergence are common at the breakpoints of old inversions ^12,13^. SVs inside the inversions showed a strong correlation with the inversion haplotype, suggesting that inversion haplotypes are evolving under strong selection.

### The evolutionary history of the inversions is marked by events of gene flux

Our phylogenomic analysis revealed that the four Atlantic herring inversions originated after the split from its sister species, the Pacific herring, between ∼1.51 and ∼2.17 MYA (Fig. 5), or 2.52 and 3.67 x 10^5^ generations ago, considering a generation time of six years for Atlantic herring ^69^. Given that ancestral *N_e_* for Atlantic herring is 4 x 10^5^ ^32^, inversions are of similar age to the coalescent time for neutral alleles (age ∼ *N_e_*), which should be enough time for exchange of variants between inversions by recombination (gene flux). In fact, our population genetics and dAF’ analyses strongly indicate the presence of gene flux in all four inversions and recombination through double crossover in the Chr6 (at ∼23.5 Mb) and Chr12 (at 23.25-24.0 Mb) inversions (Figs. 6, 8, Supplementary Figs. 7, 4A). This is also in line with previous evidence for gene flux in the Chr12 inversion ^41^. Due to gene flux, it is possible that our age estimates are underestimated.

It is expected that as inversions reach equilibrium state, gene flux erodes the divergence between haplotypes at the center but not at the breakpoints, resulting in U-shaped pattern for divergence and differentiation, as reported in some *Drosophila* species ^12,13^. Strong signals of such erosion were not observed in our data, since all four inversions showed high *F*_ST_, *d_xy_*, and strong LD (Fig. 6), which would be in line with inversions not having reached this equilibrium stage yet, despite being quite old. Similar observations are also reported in old inversions of Atlantic cod ^26^ and O_3+4_ and O_st_ inversions in *D. subobscura* ^86^. However, *F*_ST_ and LD were relatively weaker for Chr17 and 23 (Fig. 6), along with slight reduction of *d_xy_* at the center compared to the breakpoints (Supplementary Fig. 8), thus weakly supporting the expectations derived from *Drosophila* studies. The Chr6 inversion also showed high divergence at breakpoints, but the pattern continued after the distal breakpoint, which can be attributed to its presence in a high diversity region of the genome (Supplementary Fig. 8). Together, the patterns of differentiation and linkage disequilibrium among inversion haplotypes and allele sharing among Atlantic herring inversions show that gene flux contributes to the evolution of the four inversions, but it is not strong enough to completely homogenize differentiation between chromosomal arrangements given their evolutionary age, and it is probably counteracted by divergent selection for sequence polymorphisms contributing to ecological adaptation.

The nucleotide diversity shows variable patterns between derived and ancestral haplotypes across the four inversions (Fig. 6), where Chr6 and Chr12 inversions have lower diversity in the derived haplotype, as expected, while Chr17 and 23 have lower diversity in the ancestral haplotype. Higher diversity in the derived haplotype deviates from the expectations that the formation of an inversion leads to strong loss of diversity in the derived haplotype when compared to the ancestral one ^10,12,13^. However, such observation is not uncommon in natural systems ^26^ and could be explained by recovery of nucleotide diversity by the derived haplotype after the initial bottleneck when this haplotype is maintained at high frequency and in natural populations with large *N_e_*, such as Atlantic herring populations. Furthermore, gene flux between inversions could contribute to increase of nucleotide diversity of inverted haplotypes over time^13^.

### Atlantic herring inversions have evolved due to divergent selection and show no significant mutational load

The four inversions in Atlantic herring studied here show highly significant genetic differentiation among subpopulations of Atlantic herring implying a key role in local adaptation. The general pattern is that the allele named *Southern* dominates in the southern part of the species distribution while the *Northern* allele dominates in the north. It is possible that the inversions *per se* initially provided a phenotypic effect contributing to adaptation to contrasting environments in the northern and southern part of the Atlantic herring distribution, or that they captured a combination of favorable alleles at two or more loci, acting like a supergene ^11^. After their origin more than a million years ago, the alternative inversion haplotypes most likely have accumulated additional mutations contributing to fitness and that the haplotypes have been maintained by divergent selection even in the presence of gene flow between populations due to the drastic reduction in recombination caused by the inversion. Recently, divergent selection associated with local adaptation to contrasting environments has been similarly invoked to explain the maintenance of inversion polymorphisms across deer mice ^25,87^, redpolls ^24^ and Atlantic salmon ^27^.

It is generally assumed that inversion polymorphisms lead to the accumulation of genetic load due to suppression of recombination and the reduced *N_e_* for inversion haplotypes compared with other parts of the genome ^11^. Genetic load associated with inversion polymorphisms has been well documented in for instance *Drosophila* ^19^, seaweed flies ^20,88^, and *Heliconius* butterflies ^7^. However, we find no evidence for genetic load associated with the four inversions (Fig. 7). All *N* and *S* inversion haplotypes are non-lethal and found at high frequencies in different populations of this extremely abundant species which means that there must exist billions of homozygotes for each haplotype in which recombination occurs at a normal rate; given that the census population size of Atlantic herring is on the order of 10^12^ ^69^. Thus, the lack of genetic load is consistent with the presence of effective purifying selection at these loci, as we find no signature of suppressed recombination (lack of linkage disequilibrium) within inversion classes which could hamper effective purging of deleterious mutations (Supplementary Fig. 11). A similar lack of genetic load has previously been reported for other inversion polymorphisms associated with local adaptation in Atlantic cod ^26^, deer mice ^25^ and sunflower ^89^. The results suggest that accumulation of genetic load do not occur for supergenes that are fully viable in the homozygous state and when both homozygotes are common in at least some populations, as is the case for supergenes associated with local adaptation.

In conclusion, the four inversion polymorphisms characterized by long-read sequencing in this study are an important component of the arsenal of adaptive polymorphisms contributing to fitness in the Atlantic herring.

## Materials and methods

### Long-read dataset and construction of PacBio genome assemblies

Testis samples from 12 Atlantic herring, six from the Celtic Sea (collected on November 11, 2019 at latitude N51°59′ and longitude W6°48′) and six from the Baltic Sea (collected on May 18, 2020 in Hästskär, at latitude N60°35′ and longitude E17°48′) were used, representing the populations with a high frequency of the Southern (*S*) and Northern (*N*) inversion alleles ^31^, respectively. Tissue was extracted on-site and immediately flash frozen in liquid nitrogen. High molecular weight DNA was extracted using a Circulomics Nanobind Tissue Big DNA Kit (NB-900-701-001) and sized to 15-25 kb using Bioruptor (Diagenode, Denville, NJ, USA). Sequencing libraries were constructed according to the manufacturers’ protocols and each sample was sequenced on one PacBio Sequel II 8M SMRT Cell for 30 hours in circular consensus sequencing mode to generate about 20 Gb of HiFi sequence data. Similar data from an outgroup species, the European sprat (*Sprattus sprattus*), was derived from an initiative to establish a reference genome for this species (Pettersson M. E. et. al. In preparation). The quality of HiFi data for all samples was assessed using NanoPlot ^34^.

For the assembly construction, we tested two genome assemblers, HiCanu (v2.0) ^35^ and hifiasm (v0.16.1-r375) ^36^, which are specifically developed for building genome assemblies using PacBio HiFi data. Hifiasm separated the diploid genomes into primary (hap1) and secondary (hap2) haplotypes. To separate HiCanu diploid genomes, we used Purge_dups ^37^. QUAST (v5.0.2) ^38^ was used to evaluate genome statistics of all assemblies. The presence of conserved orthologs was assessed by BUSCO (v5.beta) using the vertebrate database ^39^. We noted that the secondary haplotype assemblies generated by HiCanu were more fragmented than its primary counterpart (Supplementary Table 1). Moreover, we observed that most of the breakpoint contigs from the secondary assemblies did not span the sequence around the breakpoint in one contig, hence inadequate for studying the breakpoint region. On the other hand, hifiasm arguably exceled at preserving the contiguity of all haplotypes at a phasing stage. Hence, we decided to use hifiasm assemblies for our further analyses.

### Construction of an optical genome map

The CS10 sample from the Celtic and BS3 sample from the Baltic Sea were used for optical (BioNano) mapping ^40^. Two mg of frozen testis tissue for each sample was fixed and treated according to the manufacturer’s soft tissue protocol (Bionano Genomics, San Diego, US), except that following homogenization and before fixation, the tissue suspension was passed through a 100 µm cell strainer (Miltenyi Biotec, Gaithersburg, MD). Fixed tissue was washed, and then approximately 0.7 mg was embedded in each of three agarose plugs. Embedded tissue was digested with proteinase K, treated with RNase, washed, and then equilibrated in Tris-EDTA (TE), pH 8.0. High molecular weight DNA was recovered by digesting the plugs with agarase and cleaned by a dialysis step. DNA was quantified in triplicate by Qubit (ThermoFisher) and diluted with buffer EB (Qiagen) as needed to lower the concentration to <125 ng/µL. DNA was then labeled with the DLS Labeling Kit (Bionano Genomics, San Diego, US). Recovery of labeled DNA was verified by Qubit HS dsDNA assay. Labeled molecules were linearized and imaged with the Saphyr® system (Saphyr® chip G2.3) to create the molecules data file. The single molecule image data was *de novo* assembled into optical genome maps using the hybridScaffold pipeline (Bionano Solve 3.7) with default settings (Supplementary Table 8). The assemblies were visualized using Bionano Access 1.7 webserver.

### Genome alignments of HiFi reads onto reference and PacBio assemblies

The previously reported chromosome level genome assembly ^41^ was used as a reference to align PacBio HiFi reads using minimap2 (v2.22-r1101) ^42^. Alignments with a mapping quality greater than 20 were kept using samtools ^43^ and used for further analyses. Genome-to-genome alignments were carried out using MUMmer (v4.0.0rc1) ^44,45^ with parameters “nucmer -- maxmatch -c 500 -l 200”, where all PacBio assemblies were aligned to the reference genome and to each other. The alignments for the inversion regions were visualized as dot plots using the mummerplot function of MUMmer.

### Finding inversion breakpoints using HiFi reads and constructing inversion scaffolds

In our previous study, we used PacBio Continuous long reads data from one Celtic Sea individual (CS2) to find the breakpoints for inversions on chromosomes 6 and 17 ^31^, where we visualized the alignment of a single read spanning the breakpoint using IGV ^46^ and Ribbon ^47^. Here, we used the same method to find the breakpoints on chromosomes 12 and 23 using PacBio HiFi reads and verified previously deduced breakpoints for chromosomes 6 and 17 using PacBio HiFi reads.

To compare inversion haplotypes at the sequence level, it is essential to use inversion regions in the scaffolded form. Although PacBio contigs were highly contiguous, they were not long enough to span the entire inversion regions (ranging from 1.5-8 Mb). Hence, we used optical mapping data for scaffolding PacBio contigs. However, the resulting hybrid scaffolds had many gaps and were not contiguous for the entire inversion regions (Supplementary Table 1). To overcome this, we manually curated the Bionano hybrid assemblies in the inversion regions by replacing gaps with PacBio contigs and joining hybrid scaffolds whenever necessary. We followed the NCBI recommendation to maintain a gap size of 100 (https://www.ncbi.nlm.nih.gov/genbank/wgsfaq/#q6). The correct order and orientation for PacBio contigs were decided based on their alignment to the reference assembly. The Bionano assemblies used for constructing inversion scaffolds were selected based on the contiguity of hybrid scaffolds for the respective inversion regions. As a result, we used CS10_hap1 and BS3_hap1 assemblies to make inversion scaffolds for Chr6; and CS10_hap1 and BS3_hap2 to make inversion scaffolds of Chr12, 17, and 23. This way, we had one inversion scaffold for each inversion allele for all four inversions. These scaffolds were then used for two purposes – (A) to investigate the structural variants (SVs) in the breakpoint region, and (B) as a reference to scaffold the inversion regions of the remaining 22 PacBio genomes using RagTag (v2.0.1) ^48^. The PacBio contigs were selected based on their alignment to the reference genome. The threshold for an alignment block was kept at 10 kb to avoid incorporation of non-specific contigs. However, some of the non-specific contigs had alignment blocks larger than 10 kb and had to be removed manually. The inversion scaffolds adjusted in this manner were CS7_hap1, BS2_hap1, BS4_hap2, and BS5_hap2 for the Chr17 inversion and CS4_hap2, CS5_hap2, CS7_hap1, CS10_hap2, BS5_hap1 for the Chr23 inversion. To use these scaffolds for further analysis, it was necessary to have the breakpoints of these scaffolds. However, it was challenging to apply the previously described visualization method using a single read for each scaffolded inversion because of the complexity of the breakpoint region and the presence of multiple SVs in the vicinity of the breakpoints. Hence, we used nucmer in MUMmer ^45^ to align *N* and *S* alleles from CS10_hap1 and BS3_hap2 inversion scaffolds, respectively. The resulting delta files were converted to a paf format using “delta2paf” script from paftools.js in minimap2 ^42^ to obtain the alignment co-ordinates. As only homologous sequences will align in MUMmer, the coordinates where the alignment changes its orientation would be the breakpoint. We opted for a conservative approach where SVs such as duplications, insertions, deletions, and repetitive sequences at the breakpoint regions were placed outside the inversion. This way, we first obtained breakpoints on CS10_hap1 and BS3_hap2 inversion scaffolds. They were used as a reference to obtain breakpoints from the rest of the inversion scaffolds by finding sequence homology for the 10 kb sequence near the breakpoint using BLAST (v.2.11.0+) ^49^.

### PacBio assemblies as references for *N* and *S* alleles

Although the reference genome assembly is of high quality and contiguous, it is not representative of all SVs and repeat content near the inversion breakpoints because of variation among haplotypes. Hence, we leveraged the accuracy and contiguity of HiFi assemblies and scaffolding of Bionano optical maps to build hybrid scaffolds of two assemblies (CS10_hap1 and BS3_hap2, representative of assemblies with *N* and *S* inversion alleles). We used one of each Celtic and Baltic HiFi assemblies as a reference to study structural variations, and repetitive sequences near the inversion breakpoints. As we used CS10_hap1 and BS3_hap2 assemblies to construct most of the inversion scaffolds (7 out of 8), we decided to use the same assemblies as references for structural analysis. The contigs used to build the inversion scaffolds were replaced by the inversion scaffolds in CS10_hap1 and BS3_hap2 genome assemblies. In case of the Chr6 inversion, the original inversion scaffold was built using BS3_hap1 contigs. However, the length of BS3_hap1 inversion scaffold was the same as that of BS3_hap2 (Supplementary Fig. 10) and hence, no additional modification was done for Chr6.

### Deduction of ancestral inversion allele using European sprat as an outgroup species

European sprat, an outgroup species that diverged from the Atlantic herring 11-12 MYA ^50,51^ was used to determine the ancestral inversion alleles. CS10 and BS3 inversion scaffolds, corresponding to *N* and *S* alleles, were aligned to a sprat genome assembly (Pettersson M. E. et. al. In preparation) using MUMmer ^45^. Sprat contigs corresponding to the breakpoint regions from the resulting alignment were extracted and aligned to the *N* and *S* alleles from Atlantic herring, and linear orientation of the alignment before and after the breakpoint was used to determine if the *S* or *N* allele is ancestral or derived.

### Analysis of structural variants near inversion breakpoints

To study SVs near the inversion breakpoints at the sequence level, we used one dimensional pangenome graphs and dot plots from the sequence alignments of all inversion scaffolds and a reference sequence (total of 25 sequences for each inversion). For the pangenome graph approach, we used pggb (v0.3.1) ^52^ to construct graphs and odgi (v0.7.3) ^53^ to prune the resulting graphs. To ensure that the alignments were of high-quality, we tested multiple combinations of -s (segment length) and -p (percent identity) parameters in the mapping step of pggb. We used higher -s value (20000-50000) to ensure that the graph structure represents long collinear regions of the input sequences. We used lower -p values (90-95) because inversion regions including breakpoint regions are more divergent than the rest of the genome. Exact parameters to build and visualize pangenome graphs are found on the GitHub page for this paper (https://github.com/LeifAnderssonLab/HerringInversions).

### Short-read dataset, alignment, and variant calling

The same 12 samples from Celtic and Baltic Sea used for long-read sequencing were also sequenced on Illumina HiSeq2000 sequencer to generate 2x150 bp paired-end reads of nearly 30x coverage. We assessed the read quality using FastQC 0.11.9 ^54^. We mapped reads to the reference herring genome Ch_v2.0.2 ^41^ using BWA-MEM v.0.7.17 ^55^ sorted reads with samtools v1.12^56^, and marked duplicates with Picard v2.10.3 (http://broadinstitute.github.io/picard/). To perform genotype calling for each sample, we first used Haplotyper within the Sentieon wrapper (release 201911) ^57^, which implements GATK4 HaplotypeCaller ^58^. We then combined these 12 samples with previously generated high coverage re-sequencing data for 49 Atlantic herring from Baltic Sea, Celtic Sea, North Sea, Norway, Ireland and United Kingdom, and Canada and 30 Pacific herring (*Clupea pallasii*, the sister species) distributed from the North Pacific Ocean to Norway ^31,33^. For these 91 samples, we performed joint calling of variant and invariant sites using the Genotyper algorithm within Sentieon, which implements GATK’s GenotypeGVCFs. We removed indels, and filtered genotypes with RMSMappingQuality lower than 40.0, MQRankSum lower than −12.5, ReadPosRankSum lower than −8.0, QualByDepth lower than 2.0, FisherStrand higher than 60.0 and StrandOddsRatio lower than 3.0. Additionally, we also filtered variants that had genotype quality below 20, depth below 2 or higher than three times the average coverage of the individual. These filtered vcf files were the basis for analyses therein.

### Generating consensus sequences for herring and sprat

Consensus genome sequences for each individual were generated for phylogenetic analyses. We used a custom script *do_bed.awk* ^59^ to create a bed file with the coordinates of called positions (variant and invariant sites) for each individual from the vcf files, and bedtools complement (v2.29.2) ^60^ to produce a bed file of non-called positions. We then used samtools faidx (v.1.12) and bcftools consensus (v.1.12) to introduce individual variant and invariant genotypes into the Atlantic herring reference genome, and bedtools maskfasta to hard-mask non-called positions in consensus fasta sequences.

Fasta and vcf reference sequences for the outgroup species, the European sprat, were generated by first using Chromosemble from satsuma2 (v.2016-12-07) ^61^ to align the hap1 sprat assembly to the Atlantic herring reference genome. Then, using a custom R script *ancestral_state_from_sprat.R* (https://github.com/LeifAnderssonLab/HerringInversions) that uses packages Biostrings (v.2.68.1), biomaRt (v.2.56.1), GenomicRanges (v1.52.0) and tidyverse (v.2.0.0), we extracted the regions of the sprat assembly that aligned to herring genes, choosing the longest sequence if multiple regions aligned to the same gene and excluding sprat sequences that aligned to less than 25% of the total length of genes. Then, we realigned herring and sprat sequences using mafft (v7.407) ^62^. To keep high quality alignments of true homologous regions, we further excluded alignments with missing data higher than 20% and proportion of variable sites higher than 0.2, as calculated by AMAS summary ^63^, resulting in 15,471 alignments. We converted the alignments in fasta format to a vcf file using a custom script *ancestral_vcf.py*, *genoToVcf.py*(downloaded in October 2021 from https://github.com/simonhmartin/genomics_general) and bcftools (https://github.com/LeifAnderssonLab/HerringInversions). From the final vcf file, we used the same procedure as above to generate a consensus genome sequence for the sprat in the genomic coordinates of the Atlantic herring.

### Phylogenetic inference

To obtain a maximum likelihood tree for the entire genome and for each inversion, we concatenated individual consensus genome-wide sequences of all herring individuals and the sprat. We extracted inversion alignments using samtools faidx. We removed all positions with missing data from the whole-genome alignment using AMAS trim, whereas for the inversions we allowed sites with missing data for at most 50% of the individuals. As this alignment with the sprat contained information only for 15,471 genes (114 Mb alignment, 14% of the genome), we repeated tree inference with alignments containing only herring individuals to retain more positions (346 Mb alignment, 43% of the genome) and improve the inference of intra-specific relationships, rooting the trees on the branch splitting Atlantic and Pacific herring, the typical position of sprat (Supplementary Fig. 6) ^50,51^.

### Population genomic analyses

To calculate summary statistics (differentiation as *F*_ST_, divergence as *d*_xy_, nucleotide diversity as *π*, and linkage disequilibrium as *R^2^*) within and between inverted haplotypes, we first determined the genotype of each individual in the dataset using two approaches. First, we used a set of previously ascertained highly differentiated SNPs between the *N* and *S* haplotypes at each inversion ^31,41^ and extracted genotypes for all individuals at those positions using bcftools view, keeping only positions that were polymorphic and biallelic in our dataset. The final plots were produced using the R packages tidyverse, ggplot2 (3.4.2) and ggrstar (1.0.1). Second, we performed a principal component analysis (PCA) using biallelic SNPs with less than 20% missing data and minor allele frequency (maf) above 0.01 and all individuals from the Baltic and Celtic Sea in our dataset (*N*=35) across sliding windows of 200 SNPs for chromosomes 6, 12, 17 and 23 using lostruct (downloaded October 2022 from https://github.com/petrelharp/local_pca)^64^. We genotyped individuals by plotting the first principal component for each individual across inversion regions using ggplot2 ^65^.

The genotype information obtained by these methods was further used to make four groups namely (A) all Atlantic herring (*N*=61), (B) all Pacific herring (*N*=30), (C) Number of Baltic Sea herring homozygous for *N* alleles (*N*_chr6_=20, *N*_chr12_=19, *N*_chr17_=16, *N*_chr23_=16), and (D) Number of Celtic Sea herring homozygous for *S* alleles (*N*_chr6_=15; *N*_chr12_=15, *N*_chr17_=15, *N*_chr23_=13 (Supplementary Table 9). Using a vcf file containing variant and invariant sites, we selected sites with less than 20% missing data and maf > 0.01, we calculated *d_xy_* and *F*_ST_ between these groups and *π* within groups in 20 kb sliding windows using pixy (v.1.2.5) ^66^. To study patterns of recombination suppression caused by the inversion, we calculated *R*^2^ in vcftools for both groups of homozygotes combined or individually (v.0.1.16) ^67^. For computational reasons, we used a more conservative filtering and kept sites with less than 10% missing data, genotype quality above 30 and maf above 0.1.

### Estimating the age of the inversion

We used *d_xy_* between Atlantic and Pacific herring individuals and between *N* and *S* homozygotes for each inversion to calculate the net nucleotide diversity as *d_a_* = *d_xy_* – (*d_x_*+ *d_y_*)/2 ^68^. *d_a_* was then used to estimate divergence time between Atlantic and Pacific herring, and between *N* and *S* inversion haplotypes. Assuming a mutation rate per year of *λ*=3.3x10^-10^ ^69^, we use the formula *T* = *d_a_*/2*λ* to calculate the divergence time between Atlantic and Pacific herring, and between the *N* and *S* haplotypes ^68^.

### Mutation load

To understand if recombination suppression between inverted haplotypes had resulted in the differential accumulation of deleterious mutations in inversion haplotypes, we took three main approaches. We used a similar sampling for all the analysis described above, grouping individual homozygotes for the *N* or *S* allele at each inversion to study haplotype differences. In all analyses, we used European sprat as the outgroup.

First, we calculated the ratio of number of substitutions per non-synonymous site (*d_N_*) to the number of substitutions per synonymous site (*d_S_*), or *d_N_/d_S_* between the *N* or the *S* haplotype and European sprat for all genes inside the inversions and for all genes in the genome using all 61 Atlantic herring individuals. We extracted the coding sequence of the longest isoform for each gene for each homozygote, using the consensus genome-wide sequences for each individual as described before and a combination of agat (v.0.8.0) ^70^, and bedtools getfasta. Then, using *dnds.py* (https://github.com/LeifAnderssonLab/HerringInversions) that implements biopython (v.1.79, https://biopython.org/), we calculated a consensus sequence for the *N* and *S* haplotypes using the sequences of homozygotes, converting any ambiguous positions or stop codons into missing data and removing gaps from the alignment. Finally, using alignments longer than 100 codons, we used the *cal_dn_ds* function from biopython to calculate *d_N_/d_S_* using the M0 model from codeml ^71^. We finally excluded alignments where values of *d_N_/d_S_* were higher than 3, assuming that these could be caused by alignment issues between the herring and sprat genomes. We plotted values using ggplot2 and performed a two-sided *t*-test in R 4.3.0 to test for significant differences in *d_N_/d_S_* between *N* and *S* haplotypes, and between haplotypes and the genome-wide *d_N_/d_S_* distribution.

Second, we compared the site frequency spectrum (SFS) of derived non-synonymous and synonymous mutations between *N* and *S* inversion haplotypes, using SNPEff (v5.1) ^72^ to classify the functional impact of SNPs segregating among all 61 Atlantic herring individuals. We then used a combination of bcftools and vcftools, to calculate the frequency of derived non-synonymous and synonymous biallelic SNPs (vcftools options *--freq* and --*derived*), in sites with less than 20% missing data. To run this analysis in vcftools, we used bcftools to add an extra field called *AA* to vcf files with the European sprat genotype to be used as outgroup when calculating derived allele frequencies.

Finally, we compared the proportion of transposable elements (TE) between *N* and *S* haplotypes. In this case, we used the assembled genomes for Baltic and Celtic Sea individuals and the RepeatMasker pipeline to annotate TEs. We first used RepeatModeler (2.0.1) ^73^; and the hap1 assemblies of individuals CS4, CS7, BS3 and BS4, which were either the longest and/or more contiguous of each CS and BS assemblies (Supplementary Table 1), to identify and model novel TEs. We also used BLAST (v.2.11.0+) ^74^, to compare all TEs to all protein-coding herring genes and filtered out TEs that mapped to genes. To improve the final annotation of the TEs in our database, we compared unknown repeat elements detected by RepeatModeler with transposase database (Tpases080212) ^75^ using BLAST, and used TEclassTest (v.2.1.3c) ^76^ to improve the classification of TEs in our database. We combined all four databases into one and removed redundancy with CDHit (v4.8.1) ^77^ with the parameters -c 0.9 –n 8 -d 0 -M 1600. Then, we used this library as input for RepeatMasker ^73^ to annotate and mask TEs in all the assemblies. We also annotated TEs for the inversion region of each assembly. To determine the coordinates of each inversion for each individual assembly, we used the scaffolded inversions for CS10 and BS3 (described above). We extracted 10 kb regions immediately after and before the breakpoints of the inversions from CS10 and BS3 and mapped them to the other individual assemblies using BLAST and detected the breakpoints of the inversions in each assembly with a custom script *find_breakpoints.py*that parsed the BLAST output (https://github.com/LeifAnderssonLab/HerringInversions).

### Screening for regions of recombination within inversions and enrichment of genetic variants

To further study the occurrence of gene flux between *N* and *S* inversion haplotypes, we inspected the allele frequency differences of minor alleles in *NN* and *SS* individuals combined. We first determined which allele (reference or alternative) was the minor allele in a vcf file combining *NN* and *SS* homozygotes (*N* = 35). We removed sites with minor allele frequencies below 0.2 in this combined vcf, to remove invariant sites that create noise in our dataset. Then, we (A) calculated the frequency of the minor allele in *NN* and *SS* groups separately, (B) determined the absolute difference between these frequencies, or dAF, and (C) divided dAF by the maximum frequency of the minor allele (MAFmax) in *NN* or *SS*, obtaining what we call delta allele frequency prime (dAF’). The rationale is that a new mutation occurring in *N* or *S* haplotypes will be in low frequency and will not be shared between haplotypes. As an example, consider a SNP with freq(*N*) = 0.2 and freq(*S*) = 0. dAF and dAF’ for this variant will be 0.2 and 1.0, respectively since dAF’ = abs(0.2-0.0)/0.2 = 1. If gene flux has occurred, however, the combined minor allele can be shared between inversions, resulting in lower values of dAF’ (e.g., freq(*N*) = 0.2 and freq(*S*) = 0.3, resulting in dAF’ = 0.33.

We inspected the function of variants in different dAF categories, by performing an enrichment analysis. First, we used SnpEff (v3.4) ^72^ to annotate the genome-wide variants and classify them into various categories (non-synonymous, synonymous, intronic, intergenic, 5’UTR, 3’UTR, 5 kb upstream, 5 kb downstream). Only sites within the inversion were kept for further analysis. The expected number of SNPs in each category for each inversion was calculated as p(category) X sum(extreme), where p is the total proportion of a specific SNP category without any dAF filter and sum(extreme) is the total number of SNPs with dAF > 0.95. A standard χ^2^ test was performed to test the statistical significance of the deviations of the observed values from expectation. We particularly looked at the genes that are extremely differentiated (dAF > 0.95) in the non-synonymous category.

### Conclusion

Our study sheds new light on the mechanisms that contribute to the origin and govern the evolutionary history of inversions in natural populations. Leveraging the power of long-read sequencing using multiple individuals, we deduced accurate inversion breakpoints of all four inversions and found that none of the breakpoints disrupt the coding sequence of any of the genes. We found that the majority of the inversion breakpoints are flanked by inverted duplications, possibly responsible for the origin of inversions by ectopic recombination between these sequences. The resolution provided by our population level long-read dataset also reveals that the inversion breakpoints were highly enriched for structural variants and multiple structural variants were also present within the inversions, making the inversion haplotypes highly polymorphic. Our phylogenetic and population level analyses also support that inversion polymorphisms can be maintained by divergent selection for alternatively adaptive haplotypes in the face of strong gene flow in a species with massive population sizes. We find no evidence for the accumulation of mutational load or that overdominance is important for the maintenance of inversion polymorphisms in Atlantic herring, suggesting that the high *N_e_* of *N* and *S* haplotypes combined with gene flux events should allow efficient purifying selection on both inversion alleles. Our work contributes to a better understanding of what evolutionary factors govern the maintenance of inversion polymorphisms in natural populations which is key to determine their role in adaptive evolution and speciation.

## Author contributions

LA conceived the study. MJ and MSF performed all bioinformatic analysis. EF contributed to sample collection. MEP and BWD contributed to the bioinformatic analysis. MJ, MSF, and LA wrote the paper with input from other authors. All authors approved the paper before submission.

## Data availability statement

The sequence data generated in this study is available from BioProject PRJNA1023520

## Code availability statement

The analyses of data have been carried out with publicly available software and all are cited in the Methods section. Custom scripts used are available in https://github.com/LeifAnderssonLab/HerringInversions

## Acknowledgements

We are grateful to scientists and crew on the Marine Institute’s Irish Groundfish Survey for collecting herring samples from the Celtic Sea. The study was supported by the Knut and Alice Wallenberg Foundation (KAW 2016.0361) and Vetenskapsrådet (2017-02907). The National Genomics Infrastructure (NGI)/Uppsala Genome Center provided service in massive parallel sequencing and the computational infrastructure was provided by the Swedish National Infrastructure for Computing (SNIC) at UPPMAX partially funded by the Swedish Research Council through grant agreement no. 2018-05973. M.S.F. is funded by a MSCA European Postdoctoral Fellowship (Project 101063864, INVERT2ADAPT) granted by the European Research Executive Agency.

## List of Extended data and Supplementary material

**Supplementary Fig. 1.** Dotplot of the PacBio proximal breakpoint contigs from hap1 and hap2 assemblies (Y-axis) and the reference inversion alleles (X-axis). Vertical pink lines are the inversion breakpoints and the horizontal line divides the data from the two haplotypes. **(A) Chr17** - BS5 is heterozygous for Chr17 inversion, where BS5_hap1 assembly has *S* haplotype and BS5_hap2 assembly has *N* haplotype. The homozygote (*N/N*) sample (BS1) is used as a control. **(B) Chr23** - BS2 is heterozygous for Chr23 inversion, where BS2_hap1 assembly has *N* haplotype and BS2_hap2 assembly has *S* haplotype. The homozygote sample (BS1) is used as a control with *NN* arrangement.

**Supplementary Fig. 2. (A)** Genotypes called from short read data of 91 individuals analyzed in this work at highly differentiated single nucleotide variants (SNPs) between north and south populations of Atlantic herring ^31^ and that overlap with the inversions in Chr6, Chr12, Chr17 and Chr23. Individuals are indicated in rows and positions in columns. Individuals are color coded according to their population of species of origin. Genotypes are color coded depending on their homozygosity or heterozygosity for *N* and *S* alleles. **(B)** Sliding window PCA analysis across inversion regions (sliding windows of 200 SNPs). Each line represents one of 35 individuals from the Baltic and Celtic Sea (green and yellow individuals from panel A). Individuals are color coded according to their genotype at the inversion: blue if homozygote for the *N* allele, orange if homozygote for the *S* allele and black if heterozygote. The names of heterozygote

**Supplementary Fig. 3.** Single PacBio read spanning proximal and distal breakpoint of Chr6 inversion from the CS2 sample on the reference sequence. The inversion breakpoints are shown in dotted lines.

**Supplementary Fig. 4.** Dotplot of inversion alleles. Y-axis has all inversion alleles assembled in this study from PacBio assemblies and the reference assembly. X-axis has a reference inversion *N* allele constructed using CS10_hap1 assembly. Inverted duplications at the breakpoint of Chr6, 12, and 17 inversions are indicated in the figure by black arrows above the dotplots. Red color indicates alignment in the same orientation and blue color indicates alignment in the opposite orientation.

**Supplementary Fig. 5.** Pangenome graphs using 25 sequences (24 inversion scaffolds and one reference assembly) for inversions on Chr6, 12, 17, and 23. Top 12 sequences are from Celtic Sea samples and the next 12 sequences are from Baltic Sea samples. The last sequence indicated with an asterisk is from the reference assembly. Black and red colors represent two orientations of the sequence hence, the vertical boundary of black and red are the inversion breakpoints. individuals are indicated.

**Supplementary Fig. 6.** The evolutionary history of Atlantic and Pacific herring. Maximum likelihood trees (branch lengths were ignored to facilitate visualization of relationships among individuals) that are rooted with the **(A)** European sprat or **(B)** at the branch that connects Pacific and Atlantic herring, following the topology of the tree in A. A concatenated alignment of 15471 genes (∼114 Mb) was used to produce tree in panel A, and a genome-wide alignment of 345,966,161 positions with no missing data was used to produce the tree in panel B.

**Supplementary Fig. 7.** The evolutionary history of Atlantic herring chromosomal inversions. Maximum likelihood cladograms (branch lengths of maximum likelihood trees were ignored to facilitate visualization of relationships among individuals) of concatenated alignments of chromosome 6, 12, 17 and 23 inversion regions, including the European sprat as an outgroup. Individuals are color coded by species or Atlantic herring population. Shades behind individuals indicate their inversion genotype according to Supplementary Fig. 2, and heterozygotes are indicated with a star.

**Supplementary Fig. 8.** Divergence and diversity in and around inversion regions in Atlantic herring. **(A)** Divergence (*d_xy_*) between North and South homozygotes for inversion alleles (black line) compared to divergence between Atlantic and Pacific herring (blue line). The dashed line represents the top 5% of the *d_xy_* distribution between North and South homozygotes. **(B)** Distribution of nucleotide diversity (dark gray line) of 61 Atlantic herring individuals across chromosomes 6, 12, 17 and 23 containing inversions, showing that certain inversions (e.g., chromosome 17) occur in genomic regions of elevated high nucleotide diversity.

**Supplementary Fig. 9:** Site frequency spectra of derived non-synonymous (red bars) and synonymous (blue bars) mutations for the whole genome.

**Supplementary Fig. 10.** Dotplot showing alignment of Chr6 inversion-scaffold from BS3_hap1 (Y-axis) and BS3_hap2 (X-axis).

**Supplementary Fig. 11.** Linkage disequilibrium (LD) in herring chromosomes **(A)** 6, **(B)** 12, **(C)** 17 and **(D)** 23, containing inversions. For each chromosome, LD is plotted as *R*^2^. Top plots combine Southern and Northern homozygotes (*NN*+*SS*), evidencing the high LD within the inversion regions in the context of the entire chromosome which should be recombining freely, with the exception of a few other regions of high LD. The two bottom plots only include Northern (*NN*) or Southern (*SS*) homozygotes demonstrating the lack of LD, suggesting normal recombination rate also within the inversion regions.

**Supplementary Fig. 12.** Comparison of *d_N_/d_S_* ratios for genes within the Northern (*N*) and Southern (*S*) inversion haplotypes. *d_N_/d_S_* values were calculated considering a consensus sequence for *N* and *S* haplotypes obtained from homozygotes. Genes where the ration of *d_N_/d_S_* between *N* and *S* haplotypes deviates from one are highlighted in orange. A two-sided *t*-test reveals no significant differences in the distribution of *d_N_/d_S_* values between *N* and *S* haplotypes.

**Supplementary Table 1:** Genome statistics for Celtic Sea (CS) and Baltic Sea (BS) PacBio genome assemblies where Hap1 and hap2 are the two haplotype genomes and their statistics are shown on the top and bottom row for each sample, respectively. **(A)** using hifiasm assembler **(B)** using HiCanu assembler.

**Supplementary Table 2:** Inversion breakpoint co-ordinates on the reference assembly for Chr6, Chr12, Chr17, and Chr23. *Approximate co-ordinates.

**Supplementary Table 3:** Genes within inverted region and 200 kb upstream of inversion breakpoints (shaded grey). (**A**) Chr6 inversion. (**B**) Chr12 inversion. (**C**) Chr17 inversion. (**D**) Chr23 inversion.

**Supplementary Table 4:** Inversion breakpoints for *S* (Celtic Sea) and *N* (Baltic Sea) haplotypes on four chromosomes based on individual PacBio assemblies (not the reference genome).

**Supplementary Table 5:** Structural variations surrounding inversion breakpoints.

**Supplementary Table 6:** χ^2^ test (d.f. = 1) of possible enrichment of non-synonymous mutations among extremely differentiated SNPs (dAF > 0.95).

**Supplementary Table 7:** Information on the non-synonymous SNPs showing extreme differentiation (dAF > 0.95) in the inversion regions.

**Supplementary Table 8:** Genome statistics of hybrid scaffold assemblies based on Bionano analysis. Hap1 and hap2 are the two haplotype genomes and their statistics are shown on the top and bottom row for each sample, respectively. These hybrid scaffolds included large number of Ns.

**Supplementary Table 9:** Genotypes of 35 individuals from Southern (S) and Northern (N) haplotypes at each inversion, determined by local PCA and diagnostic SNPs for each inversion.

