## Supplementary figures 1-12, supplementary tables 1, 2, 4, 5, 6, 8, 9 for "The origin and maintenance of supergenes contributing to ecological adaptation in Atlantic herring"

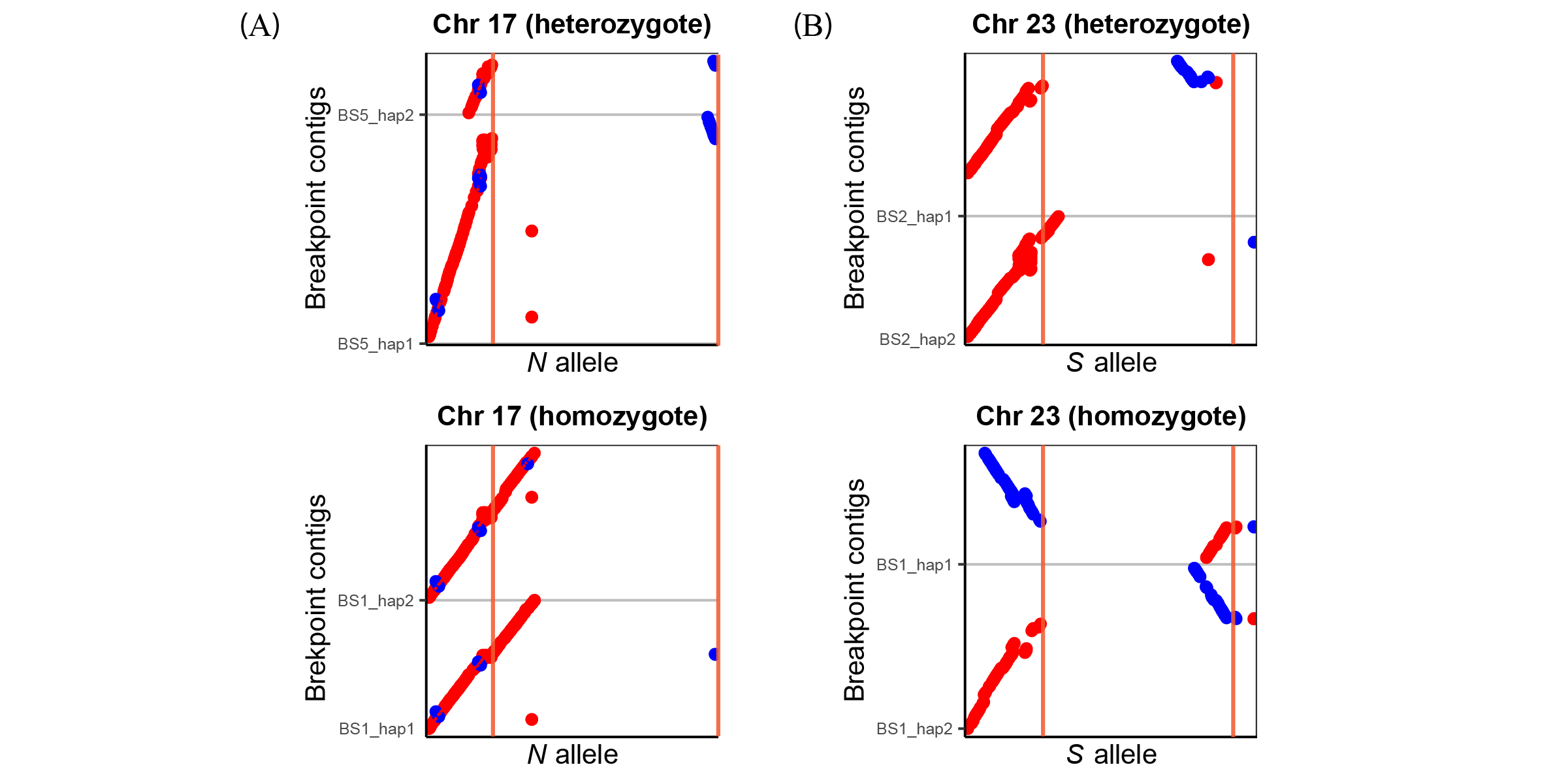

**Supplementary Fig. 1**. Dotplot of the PacBio proximal breakpoint contigs from hap1 and hap2 assemblies (Y-axis) and the reference inversion alleles (X-axis). Vertical pink lines are the inversion breakpoints and the horizontal line divides the data from the two haplotypes. **(A) Chr17** - BS5 is heterozygous for Chr17 inversion, where BS5_hap1 assembly has *S* haplotype and BS5_hap2 assembly has *N* haplotype. The homozygote (*N/N*) sample (BS1) is used as a control. **(B) Chr23** - BS2 is heterozygous for Chr23 inversion, where BS2_hap1 assembly has *N* haplotype and BS2_hap2 assembly has *S* haplotype. The homozygote sample (BS1) is used as a control with *NN* arrangement.

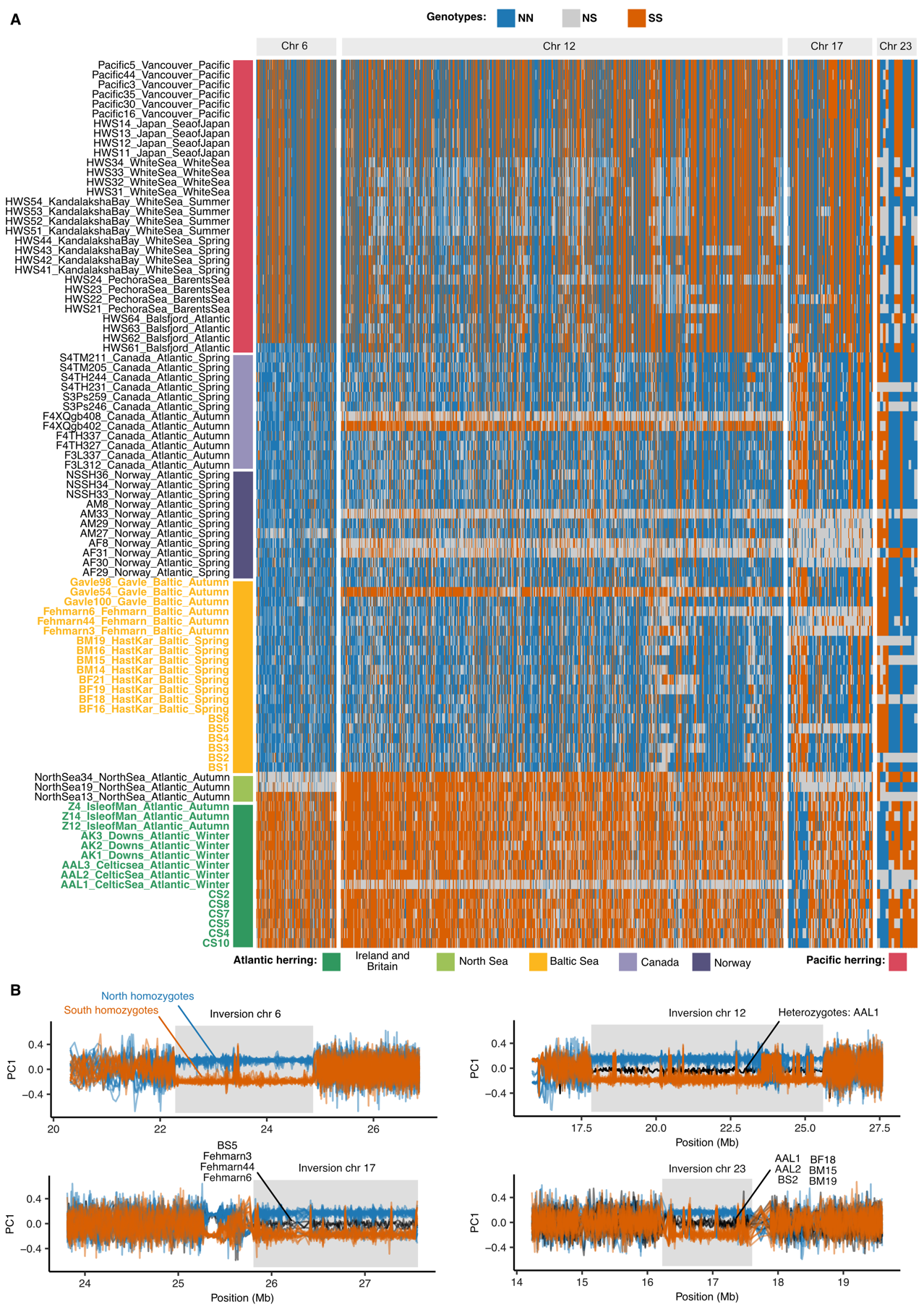

**Supplementary Fig. 2. (A)** Genotypes called from short read data of 91 individuals analyzed in this work at highly differentiated single nucleotide variants (SNPs) between north and south populations of Atlantic herring ^31^ and that overlap with the inversions on Chr6, Chr12, Chr17 and Chr23. Individuals are indicated in rows and positions in columns. Individuals are color coded according to their population of species of origin. Genotypes are color coded depending on their homozygosity or heterozygosity for *N* and *S* alleles. **(B)** Sliding window PCA analysis across inversion regions (sliding windows of 200 SNPs). Each line represents one of 35 individuals from the Baltic and Celtic Sea (green and yellow individuals from panel A). Individuals are color coded according to their genotype at the inversion: blue if homozygote for the *N* allele, orange if homozygote for the *S* allele and black if heterozygote. The names of heterozygous individuals are indicated.

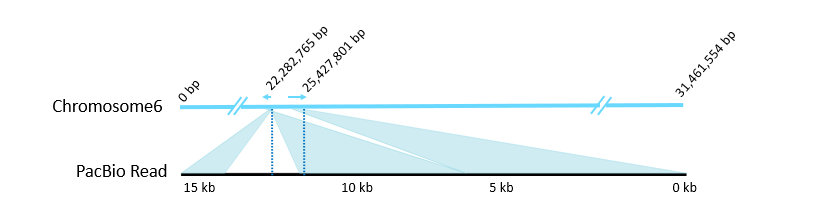

**Supplementary Fig. 3.** Single PacBio read spanning proximal and distal breakpoint of Chr6 inversion from the CS2 sample on the reference sequence. The inversion breakpoints are shown in dotted lines.

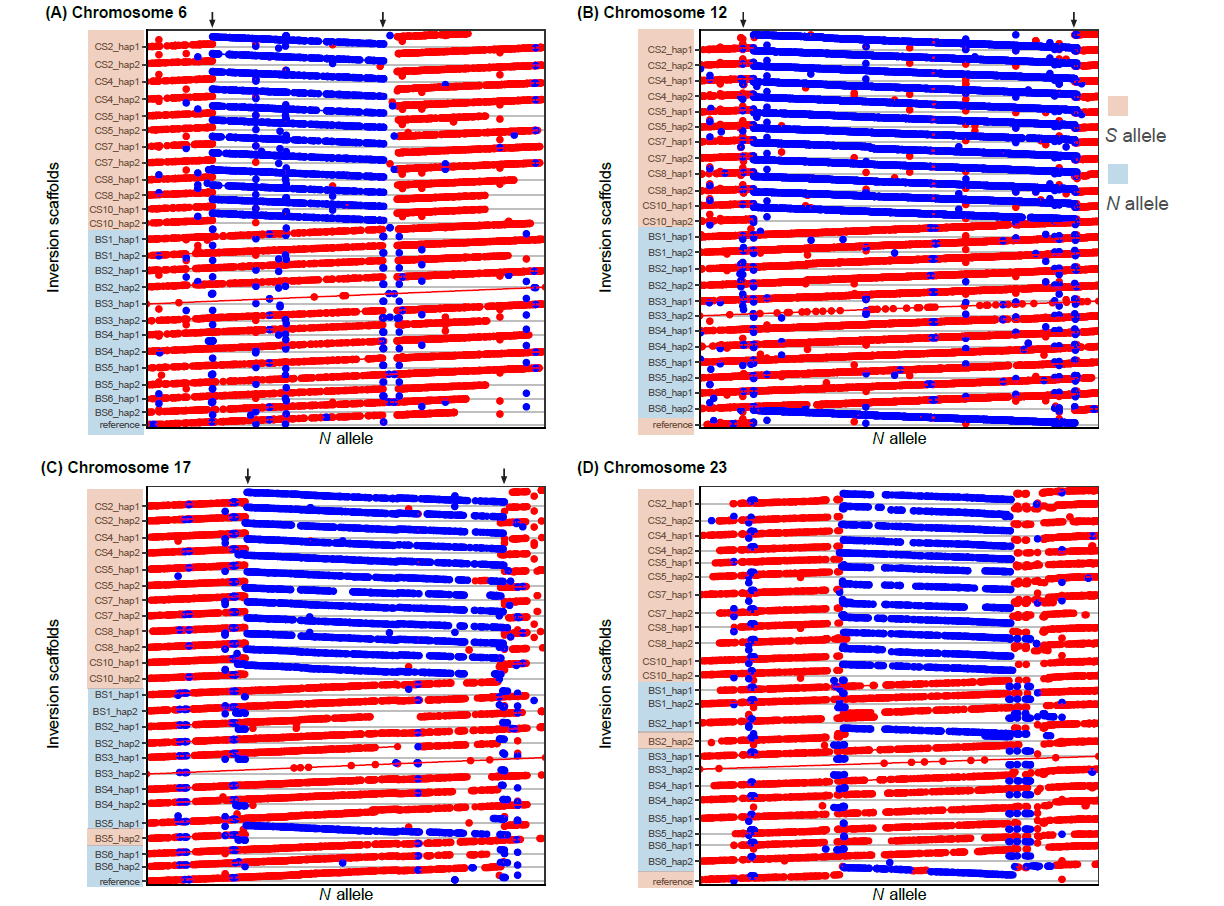

**Supplementary Fig. 4**. Dotplot of inversion alleles. Y-axis has all inversion alleles assembled in this study from PacBio assemblies and the reference assembly. X-axis has a reference inversion *N* allele constructed using CS10_hap1 assembly. Inverted duplications at the breakpoint of Chr6, 12, and 17 inversions are indicated in the figure by black arrows above the dotplots. Red color indicates alignment in the same orientation and blue color indicates alignment in the opposite orientation.

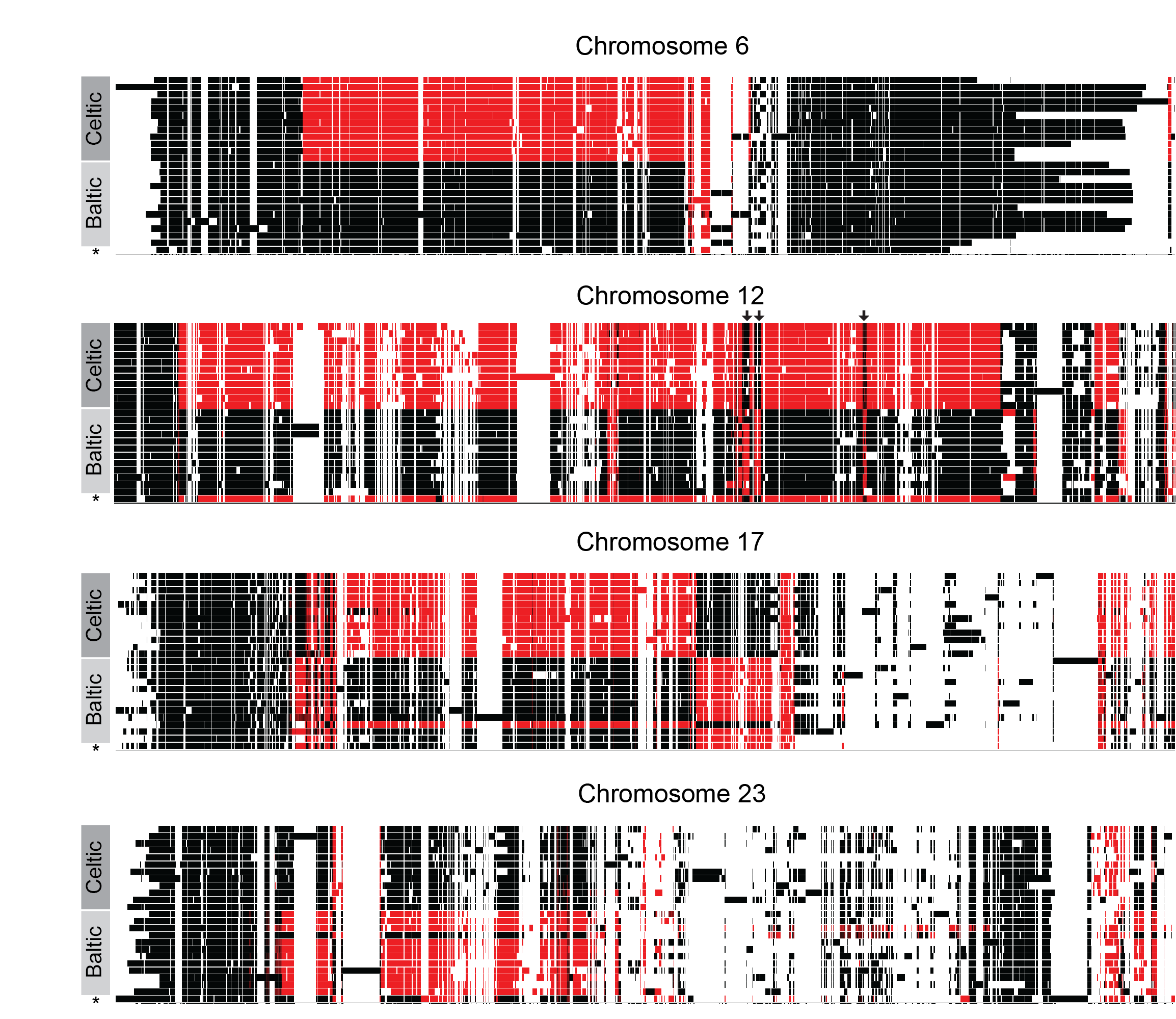

**Supplementary Fig. 5**. Pangenome graphs using 25 sequences (24 inversion scaffolds and one reference assembly) for inversions on Chr6, 12, 17, and 23. Top 12 sequences are from Celtic Sea samples and the next 12 sequences are from Baltic Sea samples. The last sequence indicated with an asterisk is from the reference assembly. Black and red colors represent two orientations of the sequence hence, the vertical boundary of black and red are the inversion breakpoints. Possible recombining regions on Chr12 inversions are indicated by black arrows at the top of its genome graph.

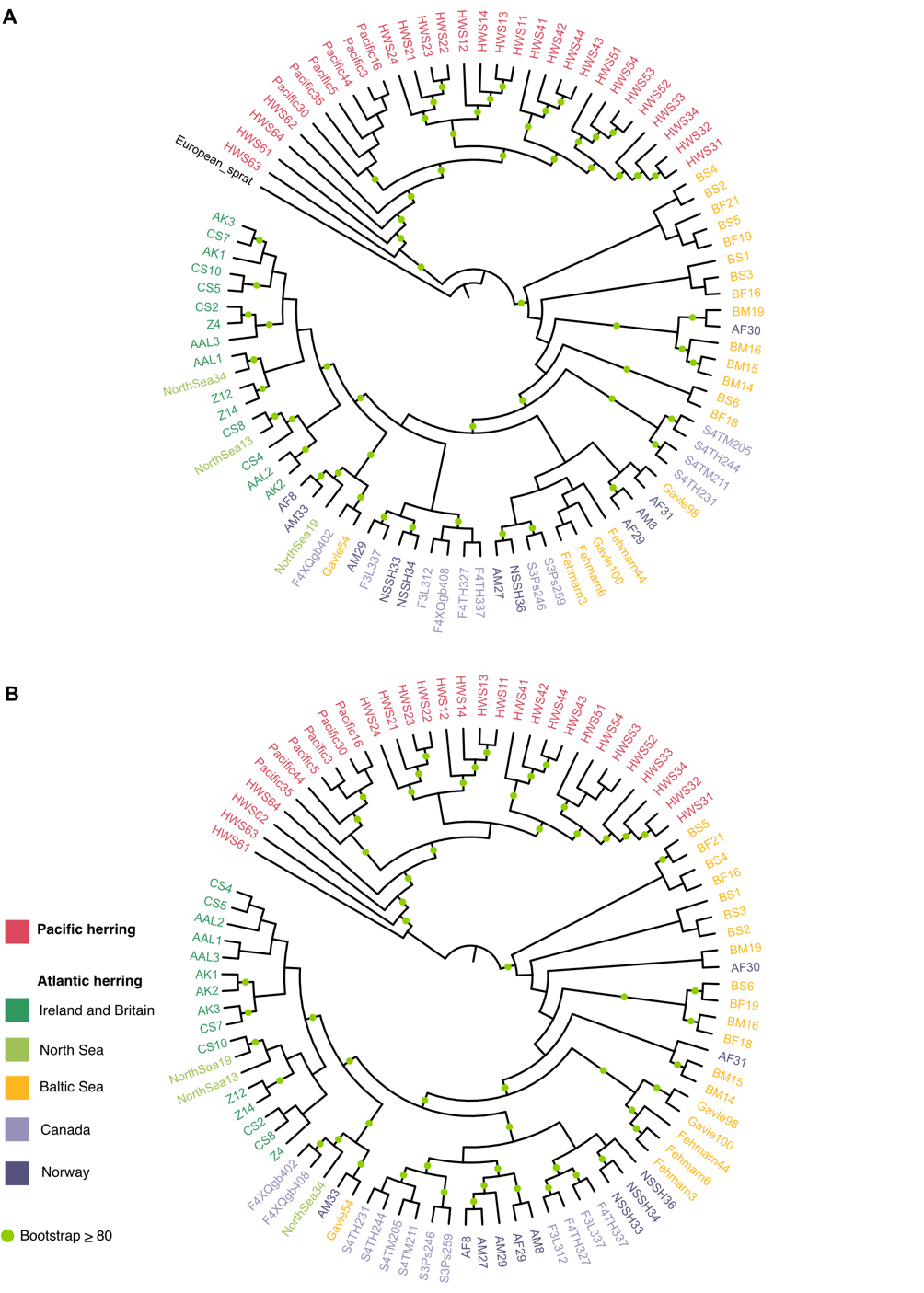

**Supplementary Fig. 6.** The evolutionary history of Atlantic and Pacific herring. Maximum likelihood trees (branch lengths were ignored to facilitate visualization of relationships among individuals) that are rooted with the **(A)** European sprat or **(B)** at the branch that connects Pacific and Atlantic herring, following the topology of the tree in A. A concatenated alignment of 15,471 genes (~114 Mb) was used to produce tree in panel A, and a genome-wide alignment of 345,966,161 positions with no missing data was used to produce the tree in panel B.

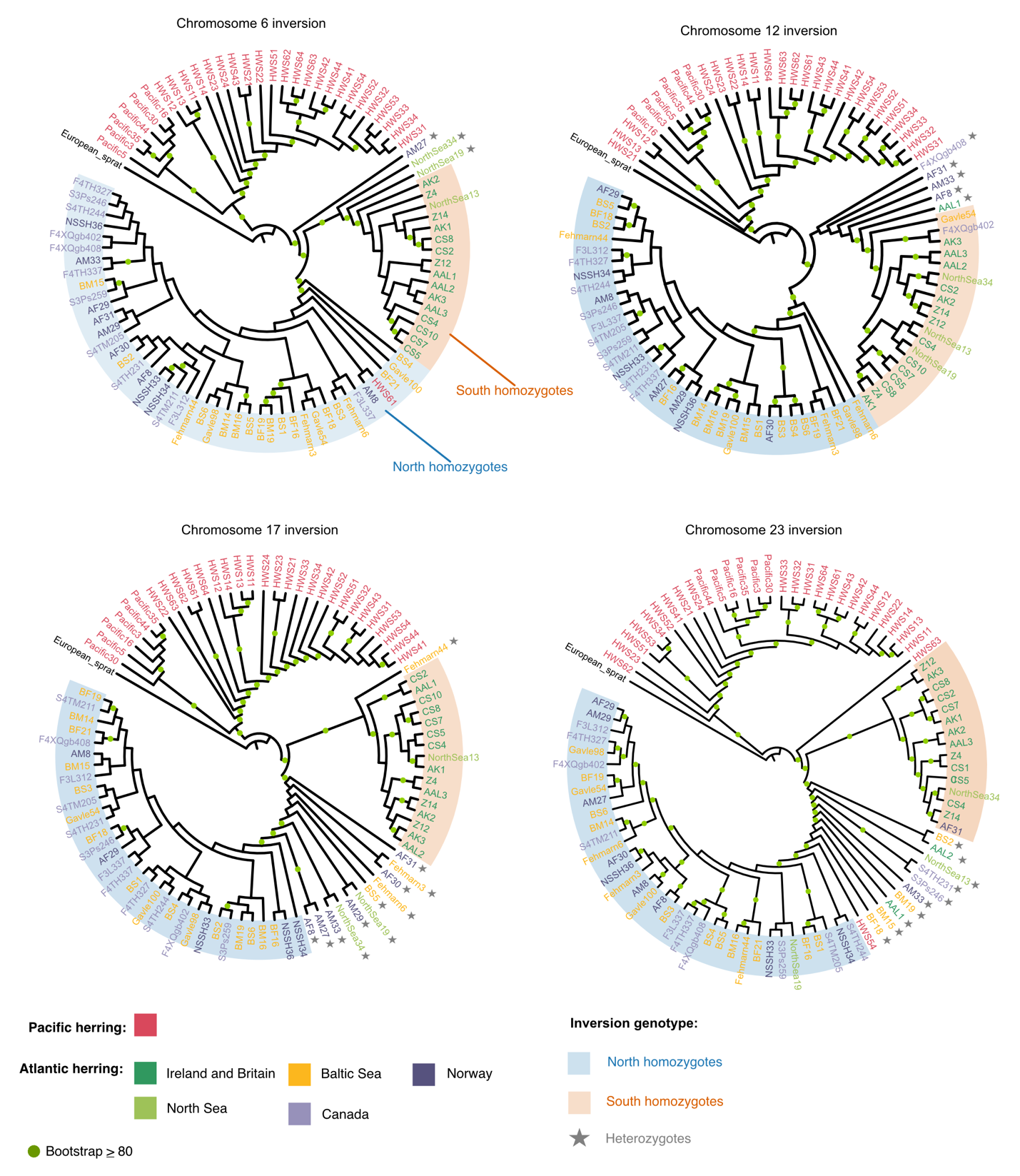

**Supplementary Fig. 7.** The evolutionary history of Atlantic herring chromosomal inversions. Maximum likelihood cladograms (branch lengths of maximum likelihood trees were ignored to facilitate visualization of relationships among individuals) of concatenated alignments of chromosome 6, 12, 17 and 23 inversion regions, including the European sprat as an outgroup. Individuals are color coded by species or Atlantic herring population. Shades behind individuals indicate their inversion genotype according to Supplementary Fig. 2, and heterozygotes are indicated with a star.

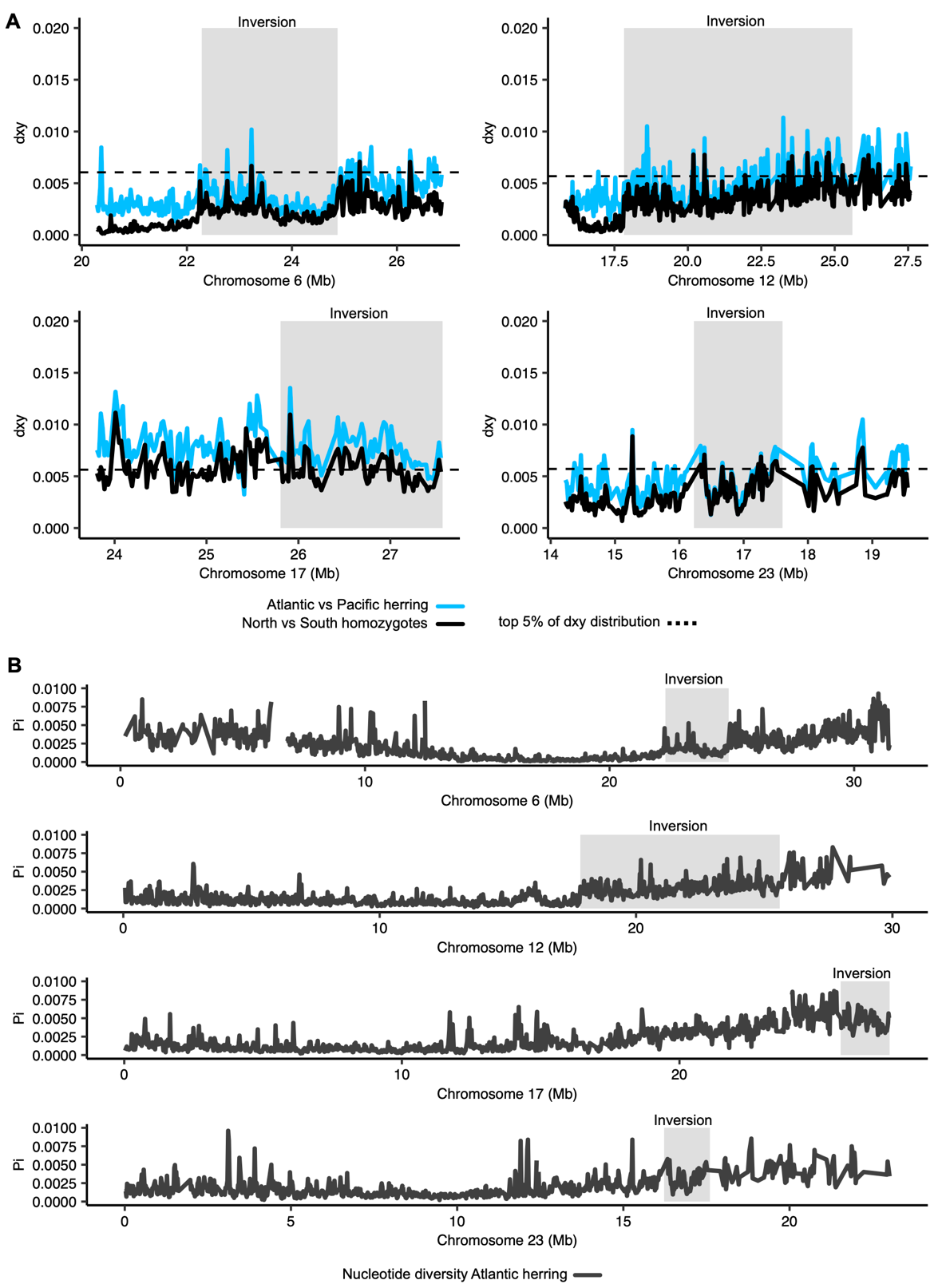

**Supplementary Fig. 8.** Divergence and diversity in and around inversion regions in Atlantic herring. **(A)** Divergence (*d_xy_*) between North and South homozygotes for inversion alleles (black line) compared to divergence between Atlantic and Pacific herring (blue line). The dashed line represents the top 5% of the *dxy* distribution between North and South homozygotes. **(B)** Distribution of nucleotide diversity (dark gray line) of 61 Atlantic herring individuals across chromosomes 6, 12, 17 and 23 containing inversions, showing that certain inversions (e.g., chromosome 17) occur in genomic regions of elevated high nucleotide diversity.

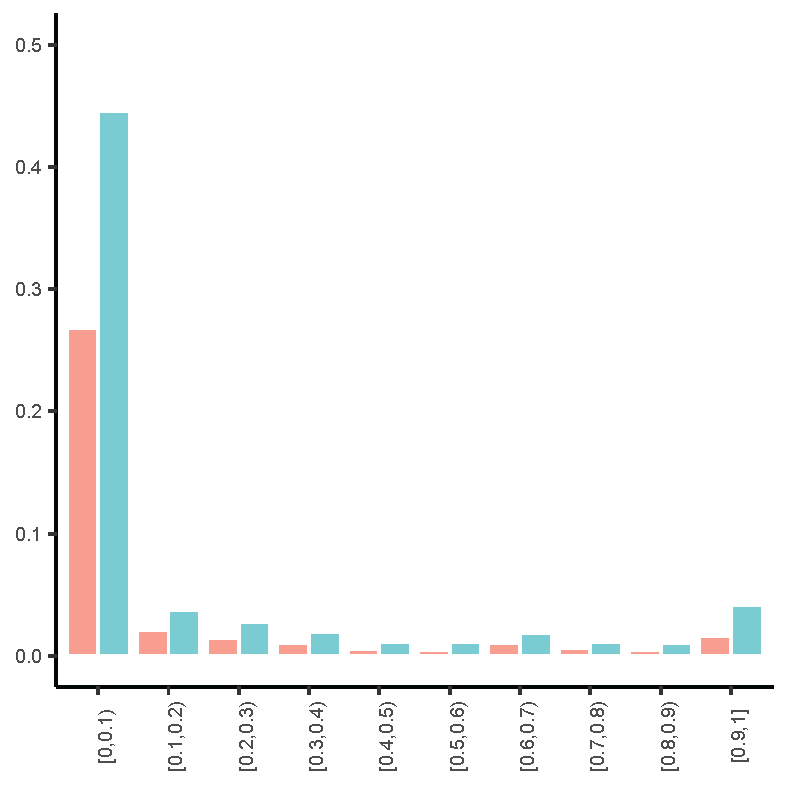

**Supplementary Fig. 9:** Site frequency spectra of derived non-synonymous (red bars) and synonymous (blue bars) mutations for the whole genome.

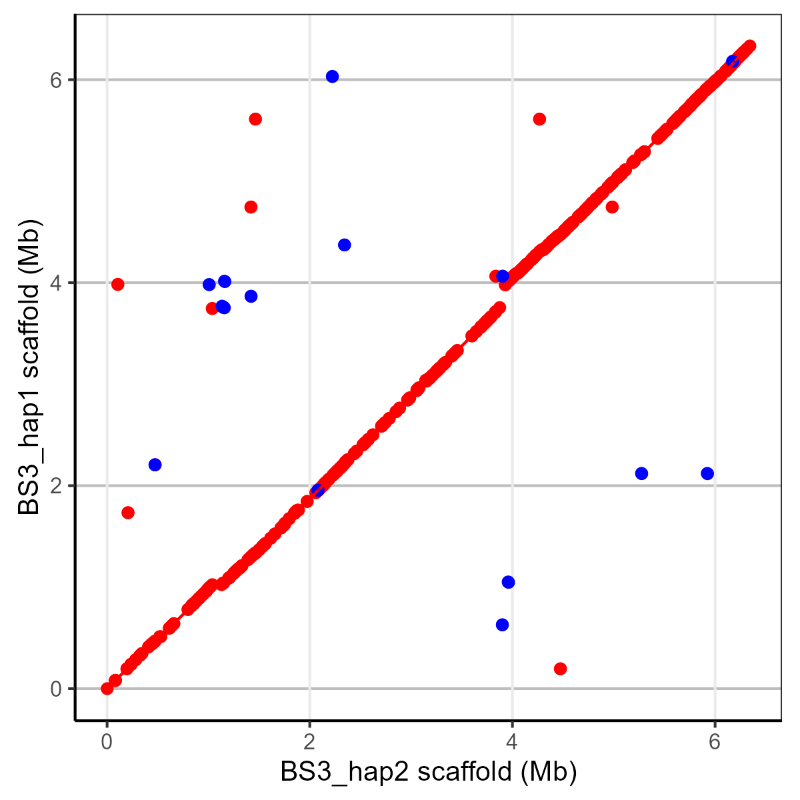

**Supplementary Fig. 10**. Dotplot showing alignment of Chr6 inversion-scaffold from BS3_hap1 (Y-axis) and BS3_hap2 (X-axis).

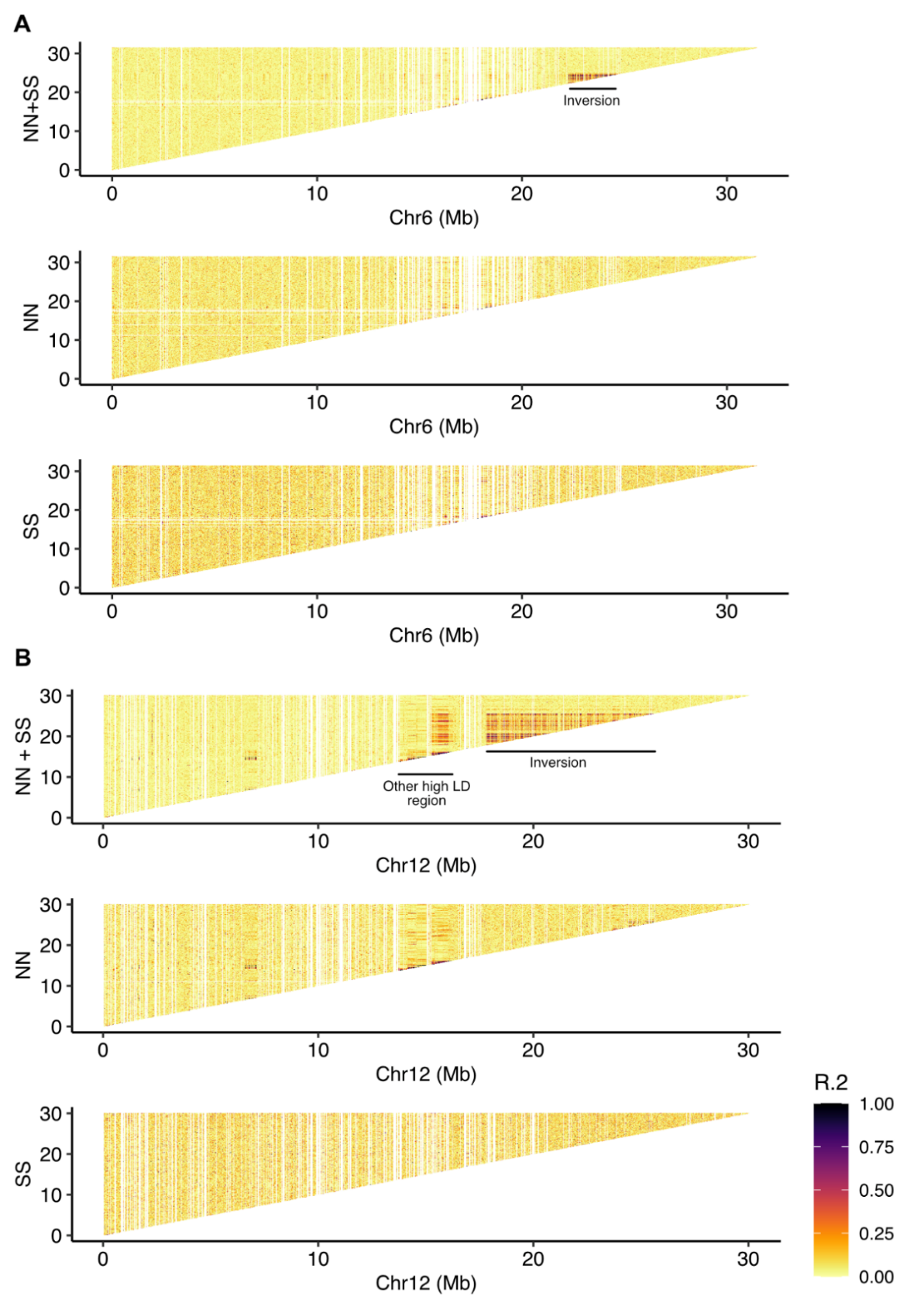

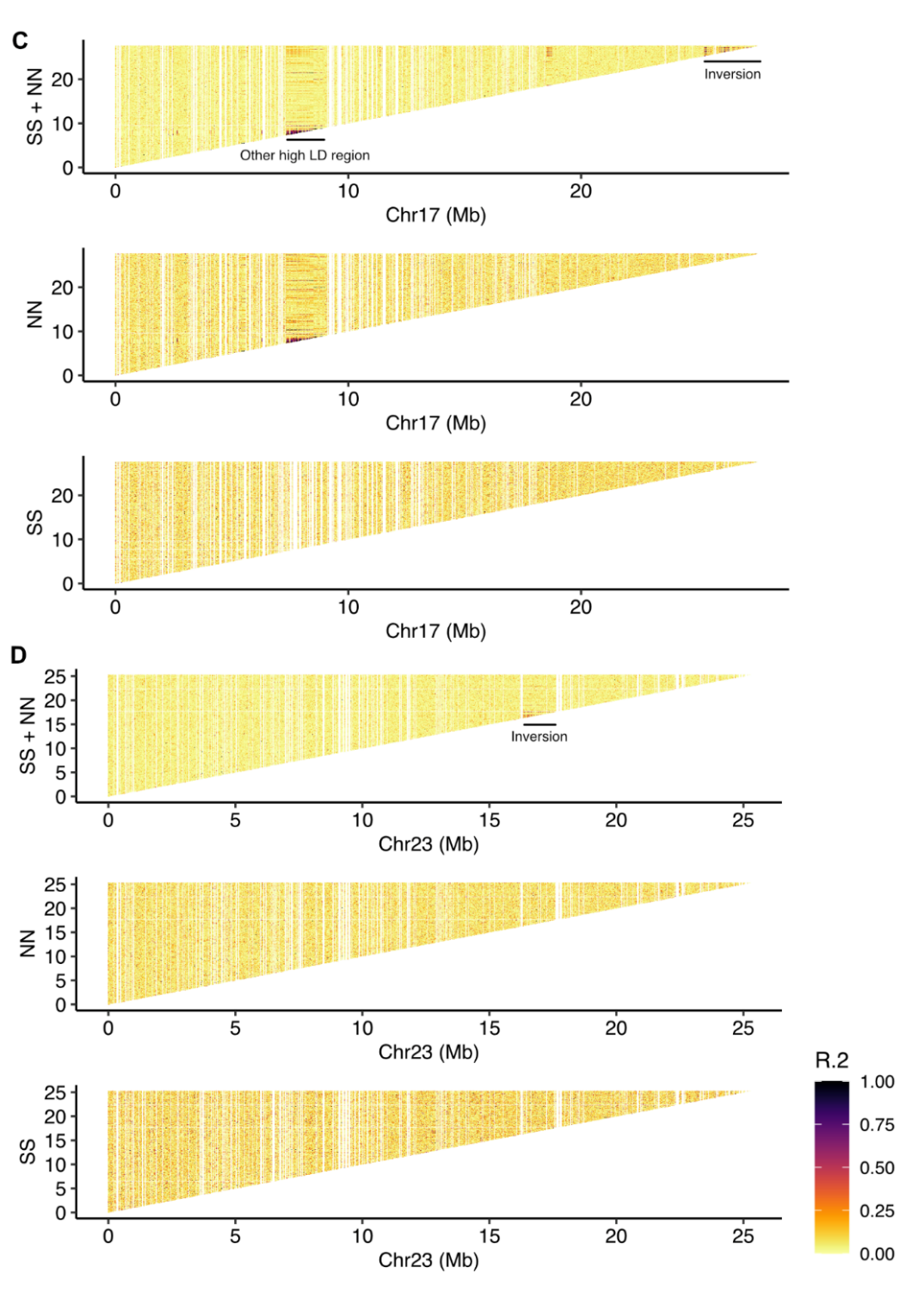

**Supplementary Fig. 11.** Linkage disequilibrium (LD) in herring chromosomes **(A)** 6, **(B)** 12, **(C)** 17 and **(D)** 23, containing inversions. For each chromosome, LD is plotted as *R^2^*. Top plots combine Southern and Northern homozygotes (*NN*+*SS*), evidencing the high LD within the inversion regions in the context of the entire chromosome which should be recombining freely, with the exception of a few other regions of high LD. The two bottom plots only include Northern (*NN*) or Southern (*SS*) homozygotes demonstrating the lack of LD, suggesting normal recombination rate also within the inversion regions.

**Supplementary Fig. 12.** Comparison of *d_N_/d_S_* ratios for genes within the Northern (*N*) and Southern (*S*) inversion haplotypes. *d_N_/d_S_* values were calculated considering a consensus sequence for *N* and *S* haplotypes obtained from homozygotes. Genes where the ration of *d_N_/d_S_* between *N* and *S* haplotypes deviates from one are highlighted in orange. A two-sided *t*-test reveals no significant differences in the distribution of *d_N_/d_S_* values between *N* and *S* haplotypes.

| **Samples** | **Genome size (Mb)** | **Genome completeness (%)** | **No. contigs** | **N50 (Kb)** |
| --- | --- | --- | --- | --- |
| **CS2** | 769.1 | 91.5 | 4364 | 524.8 |
|  | 744.6 | 91.6 | 3314 | 547.9 |
| **CS4** | 776.3 | 92.8 | 3253 | 693.1 |
|  | 751.3 | 92.8 | 2539 | 731.1 |
| **CS5** | 768.1 | 91.6 | 3817 | 545.0 |
|  | 745.9 | 91.6 | 3037 | 569.8 |
| **CS7** | 783.6 | 92.0 | 3929 | 469.5 |
|  | 751.1 | 90.6 | 3117 | 512.3 |
| **CS8** | 781.7 | 91.9 | 4100 | 526.5 |
|  | 743.2 | 91.5 | 3280 | 504.5 |
| **CS10** | 773.1 | 90.7 | 3997 | 452.8 |
|  | 756.6 | 90.3 | 3228 | 472.4 |
| **BS1** | 776.2 | 91.6 | 3756 | 576.2 |
|  | 751.8 | 91.4 | 3028 | 606.5 |
| **BS2** | 788.3 | 91.8 | 3769 | 540.9 |
|  | 759.7 | 90.7 | 2952 | 539.8 |
| **BS3** | 779.2 | 92.4 | 3331 | 680.2 |
|  | 758.5 | 92.1 | 2725 | 661.8 |
| **BS4** | 792.2 | 92.8 | 3309 | 737.1 |
|  | 760.1 | 92.0 | 2570 | 735.4 |
| **BS5** | 780.3 | 90.6 | 4045 | 494.0 |
|  | 751.8 | 91.1 | 3259 | 480.0 |
| **BS6** | 787.8 | 92.2 | 3771 | 572.2 |
|  | 764.3 | 92.3 | 3083 | 557.1 |
| **European sprat** | 850.3 | 91.8 | 4748 | 544.6 |
|  | 802.4 | 91.7 | 3611 | 590.2 |
| **Supplementary Table 1A**: Genome statistics for Celtic Sea (CS) and Baltic Sea (BS) PacBio genome assemblies using hifiasm assembler. Hap1 and hap2 are the two haplotype genomes and their statistics are shown on the top and bottom row for each sample, respectively. | | | | |

| **Samples** | **Genome size (Mb)** | **No. contigs** | **N50 (Kb)** |
| --- | --- | --- | --- |
| **CS2** | 747.7 | 3497 | 554.3 |
|  | 774.7 | 9155 | 234.1 |
| **CS4** | 797.7 | 3029 | 744.8 |
|  | 721.9 | 5289 | 306.9 |
| **CS5** | 760.0 | 3202 | 564.2 |
|  | 728.4 | 6000 | 267.7 |
| **CS7** | 759.4 | 3468 | 489.6 |
|  | 710.3 | 5736 | 244.7 |
| **CS8** | 760.6 | 3347 | 553.2 |
|  | 738.0 | 6330 | 243.4 |
| **CS10** | 763.6 | 3447 | 507.1 |
|  | 710.8 | 5969 | 231.9 |
| **BS1** | 626.3 | 3064 | 626.3 |
|  | 281.3 | 5584 | 281.3 |
| **BS2** | 586.6 | 3000 | 586.6 |
|  | 265.6 | 5802 | 265.6 |
| **BS3** | 724.0 | 2682 | 724.0 |
|  | 321.1 | 5559 | 321.1 |
| **BS4** | 805.2 | 2615 | 805.2 |
|  | 342.4 | 5750 | 342.4 |
| **BS5** | 531.4 | 3357 | 531.4 |
|  | 241.5 | 5983 | 241.5 |
| **BS6** | 606.3 | 3083 | 606.3 |
|  | 270.9 | 5797 | 270.9 |
| **Supplementary Table 1B**: Genome statistics for Celtic Sea (CS) and Baltic Sea (BS) PacBio genome assemblies using HiCanu assembler. Hap1 and hap2 are the two haplotype genomes and their statistics are shown on the top and bottom row for each sample, respectively. | | | |

| **Inversion** | **Samples** | **Proximal breakpoint** | **Distal breakpoint** |
| --- | --- | --- | --- |
| Chr6 | CS2 | 22,282,765 | 25,427,801 |
| Chr6 | CS4 | 22,282,765 | 24,869,682 |
| Chr6 | CS7 | 22,282,765 | 24,869,682 |
| Chr6 | CS10 | 22,282,765 | 24,869,682 |
| Chr12 | BS1 | 17,818,247 | 25,598,057* |
| Chr12 | BS2 | 17834922* | 25,597,304* |
| Chr12 | BS3 | 17840422* | 25,603,093* |
| Chr12 | BS4 | 17826319* | 25,603,093 |
| Chr12 | BS5 | 17841789* | 25,603,093* |
| Chr12 | BS6 | 17826319* | 25,603,093* |
| Chr17 | CS2 | 25,802,209 | 27,568,510 |
| Chr17 | CS4 | 25,802,209 | 27,568,510 |
| Chr17 | CS5 | 25,802,212 | 27,568,510 |
| Chr17 | CS7 | 25,802,212 | 27,568,510 |
| Chr17 | CS8 | 25,802,209 | 27,568,510 |
| Chr17 | CS10 | 25,802,209 | 27,568,510 |
| Chr23 | BS1 | 16,225,343 | 17,604,279 |
| Chr23 | BS2 | 16,216,922 | 17,604,291 |
| Chr23 | BS3 | 16,225,343 | 17,604,291 |
| Chr23 | BS4 | 16,225,343 | 17,604,279 |
| Chr23 | BS5 | 16,225,343 | 17,604,277 |
| Chr23 | BS6 | 16,226,443 | 17,603,173 |
| **Supplementary Table 2**: Inversion breakpoint co-ordinates on the reference assembly for Chr6, Chr12, Chr17, and Chr23. *Approximate co-ordinates. | | | |

| **Inversion** | **Sample** | **Proximal breakpoint** | **Distal breakpoint** |
| --- | --- | --- | --- |
| **Chr6** | **Celtic** | 1144464 | 3774578 |
| **Chr6** | **Baltic** | 1051601 | 3747148 |
| **Chr12** | **Celtic** | 1156916 | 8338930 |
| **Chr12** | **Baltic** | 1215813 | 8428837 |
| **Chr17** | **Celtic** | 741635 | 2249708 |
| **Chr17** | **Baltic** | 687774 | 2468994 |
| **Chr23** | **Celtic** | 854555 | 2134906 |
| **Chr23** | **Baltic** | 788123 | 1676584 |
| **Supplementary Table 4**: Inversion breakpoints for *S* (Celtic Sea) and *N* (Baltic Sea) haplotypes on four chromosomes based on individual PacBio assemblies (not the reference genome). | | | |

| **Inversion** | **Location** | **Type of SV** | **Description** | **Gene** |
| --- | --- | --- | --- | --- |
| Chr6 | Proximal and distal | Inverted duplication of 35 kb in S alleles and 45 kb in N alleles | In the reference, co-ordinates are - chr6:22223000-22261500 and chr6:24913000-24951000 | BTLN-2 |
|  | Proximal | 30-50 kb insertion in CS10_hap1, CS10_hap2, BS2_hap2, BS3_hap2, BS6_hap1 |  |  |
|  | Distal | 30-200 kb divergent sequence. 200 kb sequence is present in BS3_hap1, BS5_hap2, and BS6_hap2 |  |  |
| Chr12 | Proximal and distal | Inverted duplication of 8 kb in both S and N alleles | In the reference, co-ordinates are chr12:17818247-17826319 and chr12:25602941-25610712 | FUT-9 |
|  | Distal | 30 kb palindromic sequence |  |  |
| Chr17 | Proximal and distal | Inverted duplication of 60 kb in both S and N alleles | In the reference, co-ordinates are chr17:25728061-25805444 | NA |
|  | Proximal | 20 kb sequence in one-three copies in all S and N alleles |  | fatty acid binding protein, liver-type-like |
|  | Distal | 50 kb palindromic sequence in all S and N alleles except BS1_hap1, BS3_hap1, BS4_hap2, BS6_hap2. These exception alleles also don't have a telomeric sequence outside inversion. BS5_hap2 has one more palindromic sequence. It is absent in the reference assembly. | One of the copies is very fragmented. Has many repeats. | NA |
|  |  | The telomeric sequence next to the breakpoint is of variable length in all assemblies, ranging from 0 kb to 300 kb |  |  |
| Chr23 | Proximal | 130 kb sequence is present in 3-4 copies in all S and N alleles | Additional copy of this sequence is present in the opposite orienation in all N alleles, except BS4_hap2. It is absent in all S alleles except CS2_hap2 and CS10_hap1. | NA |
|  |  | 13 kb insertion in eight N alleles; BS2_hap1, BS2_hap2, BS3_hap1, BS3_hap2, BS4_hap1, BS4_hap2, BS5_hap1, BS6_hap2. |  | NACHT, LRR and PYD domains-containing protein 12-like |
|  | Distal | 20 kb sequence in two copies. Only 10 kb of it is present in all S and N alleles. The entire 20 kb is present in CS7_hap2, CS8_hap1, and BS2_hap1 |  | nuclear body protein SP140-like protein |
|  |  | Insertion of variable lengths ranging from 50 to 700 kb. (CS2_hap2 at proximal, 300 kb CS5_hap1 at distal, 50 kb; CS5_hap2 at distal, 100 kb; CS7_hap1 and hap2 at distal, 700 kb; CS8_hap1 at distal, 200 kb; CS8_hap2 at distal, 300 kb, 150 kb of it is homologous to that of CS8_hap1; CS10_hap1 at distal, 250 kb; BS1_hap1 at distal, around 250 kb, fragmented BS3_hap1 at distal, 100 kb; BS5_hap1 at distal, 500 kb) |  |  |
| **Supplementary Table 5**: Structural variations surrounding inversion breakpoints | | | | |

| **Inversion** | **Observed** | **Expected** | ***P*-value (d.f. = 1)** |
| --- | --- | --- | --- |
| Chr6 | 7 | 5 | 0.3677 |
| Chr12 | 13 | 9 | 0.1805 |
| Chr17 | 2 | 2 | 1.0000 |
| Chr23 | 3 | 2 | 0.4737 |
| **Supplementary Table 6**: *χ*^2^ test (d.f. = 1) of possible enrichment of non-synonymous mutations among extremely differentiated SNPs (dAF > 0.95). | | | |

| **Samples** | **Genome size (Mb)** | **No. contigs** | **N50 (Mb)** | **N's per 100 kb (Kb)** | **Not scaffolded sequence (Mb)** |
| --- | --- | --- | --- | --- | --- |
| **CS10** | 937.3 | 123 | 23.0 | 28.6 | 104.2 |
|  | 950.5 | 129 | 22.9 | 29.3 | 85.0 |
| **BS3** | 866.7 | 129 | 23.5 | 19.9 | 85.1 |
|  | 871.4 | 135 | 23.4 | 21.0 | 70.0 |
| **Supplementary Table 8**: Genome statistics of hybrid scaffold assemblies based on Bionano analysis. Hap1 and hap2 are the two haplotype genomes and their statistics are shown on the top and bottom row for each sample, respectively. These hybrid scaffolds included large number of Ns. | | | | | |

| **Individual** | **Population** | **Inversion** | | | |
| --- | --- | --- | --- | --- | --- |
|  |  | **Chr6** | **Chr12** | **Chr17** | **Chr23** |
| AAL1_CelticSea_Atlantic_Winter | Celtic Sea | SS | NS | SS | NS |
| AAL2_CelticSea_Atlantic_Winter | Celtic Sea | SS | SS | SS | NS |
| AAL3_Celticsea_Atlantic_Winter | Celtic Sea | SS | SS | SS | SS |
| CS10 | Celtic Sea | SS | SS | SS | SS |
| CS4 | Celtic Sea | SS | SS | SS | SS |
| CS5 | Celtic Sea | SS | SS | SS | SS |
| CS7 | Celtic Sea | SS | SS | SS | SS |
| CS8 | Celtic Sea | SS | SS | SS | SS |
| CS2 | Celtic Sea | SS | SS | SS | SS |
| AK1_Downs_Atlantic_Winter | Celtic Sea | SS | SS | SS | SS |
| AK2_Downs_Atlantic_Winter | Celtic Sea | SS | SS | SS | SS |
| AK3_Downs_Atlantic_Winter | Celtic Sea | SS | SS | SS | SS |
| Z12_IsleofMan_Atlantic_Autumn | Celtic Sea | SS | SS | SS | SS |
| Z14_IsleofMan_Atlantic_Autumn | Celtic Sea | SS | SS | SS | SS |
| Z4_IsleofMan_Atlantic_Autumn | Celtic Sea | SS | SS | SS | SS |
| BS1 | Baltic Sea | NN | NN | NN | NN |
| BS2 | Baltic Sea | NN | NN | NN | NS |
| BS3 | Baltic Sea | NN | NN | NN | NN |
| BS4 | Baltic Sea | NN | NN | NN | NN |
| BS5 | Baltic Sea | NN | NN | NS | NN |
| BS6 | Baltic Sea | NN | NN | NN | NN |
| Fehmarn3_Fehmarn_Baltic_Autumn | Baltic Sea | NN | NN | NS | NN |
| Fehmarn44_Fehmarn_Baltic_Autumn | Baltic Sea | NN | NN | NS | NN |
| Fehmarn6_Fehmarn_Baltic_Autumn | Baltic Sea | NN | NN | NS | NN |
| Gavle100_Gavle_Baltic_Autumn | Baltic Sea | NN | NN | NN | NN |
| Gavle54_Gavle_Baltic_Autumn | Baltic Sea | NN | SS | NN | NN |
| Gavle98_Gavle_Baltic_Autumn | Baltic Sea | NN | NN | NN | NN |
| BF16_HastKar_Baltic_Spring | Baltic Sea | NN | NN | NN | NN |
| BF18_HastKar_Baltic_Spring | Baltic Sea | NN | NN | NN | NS |
| BF19_HastKar_Baltic_Spring | Baltic Sea | NN | NN | NN | NN |
| BF21_HastKar_Baltic_Spring | Baltic Sea | NN | NN | NN | NN |
| BM14_HastKar_Baltic_Spring | Baltic Sea | NN | NN | NN | NN |
| BM15_HastKar_Baltic_Spring | Baltic Sea | NN | NN | NN | NS |
| BM16_HastKar_Baltic_Spring | Baltic Sea | NN | NN | NN | NN |
| BM19_HastKar_Baltic_Spring | Baltic Sea | NN | NN | NN | NS |
| **Supplementary Table 9:** Genotypes of 35 individuals from Southern (*S*) and Northern (*N*) haplotypes at each inversion, determined by local PCA and diagnostic SNPs for each inversion. | | | | | |
